# Single-cell multiplex approaches deeply map ON-target CRISPR-genotoxicity and reveal its mitigation by palbociclib and long-term engraftment

**DOI:** 10.1101/2025.04.04.647179

**Authors:** Julian Boutin, Sabrina Fayet, Victor Marin, Camille Bergès, Maude Riandière, Jérôme Toutain, Isabelle Lamrissi-Garcia, Chloé Thibault, David Cappellen, Sandrine Dabernat, Arthur Poulet, Maëla Francillette, Nathalie Droin, Christelle Debeissat, Philippe Brunet de la Grange, François Moreau-Gaudry, Aurélie Bedel

## Abstract

Genome editing by CRISPR-Cas9 is promising for gene therapy. However, safety concerns remain, particularly regarding the ON-target genotoxicity associated with protocols using nucleases. Monitoring the genotoxicity of edited cells before and after graft is essential, especially to assay potentially deleterious megabase-scale genomic rearrangements induced at the targeted *locus*. High sensitivity requires single-cell resolution. Here, we developed an integrated approach combining targeted single-cell DNA sequencing focused on single nucleotide polymorphism (scSNP-DNAseq) with complementary micronuclei and LOH cytometry-reporter assays. This multiplexed strategy enables orthogonal readouts to accurately monitor CRISPR-mediated genotoxicity in primary cells. Using this approach, we detected, mapped and characterized various types of induced-losses of heterozygosity (terminal, interstitial, copy-loss and copy-neutral). Our compelling workflow assessed editing-associated chromosomal instability linked to double strand break after editing. Importantly, palbociclib prevented the appearance of such genomic rearrangements in hematopoietic stem/progenitor cells without impairing cell fate or graft capability. Conversely, short-term risk was significantly increased with DNA-PKcs inhibitor AZD7648 (HDR booster) in HSPCs and fibroblasts. Fortunately, targeting *<u>HBG1/2</u>* in Chr11p <u>in HSPC</u>s, scSNP-DNA-seq revealed that ON-target genotoxic events were no longer detectable after long-term xenografts, even in AZD7648-treated cells. This work demonstrates that scSNP-DNA-seq should be routinely implemented to monitor chromosomal rearrangements before and after CRISPR-edited cell infusions.

## Introduction

CRISPR-Cas9 genome editing holds tremendous promise for gene therapy^1–4^. It offers unprecedented potential for therapeutic applications. Numerous CRISPR-based clinical trials are ongoing and the first treatments are already approved for clinical use. Despite development of new genome editing tools, most of current clinical applications still rely on CRISPR-Cas9 nuclease, and create a targeted DNA double-strand break (DSB). However, the repair of DSB can lead to complex and unpredictable outcomes, raising concerns about genotoxic risk and potential phenotypic alterations in edited cells. Both OFF-target and ON-target genotoxicity have been reported. ON-target genotoxicity includes various types of loss-of-heterozygosities (LOH): kilobase-scale interstitial copy-loss by deletions^5^ (CL-LOH), megabase-scale telomeric CL-LOH (truncations)^6^, and copy-neutral loss-of-heterozygosity (CN-LOH) which may lead to imprinting loss and disrupt imprinted genes expression^7^. Even larger genomic abnormalities have been observed, such as whole-chromosome loss^8^ and chromothripsis^9–11^. Importantly, these genomic rearrangements are observed *in vitro,* a few days after editing, and have been observed in many cell types, in immortalized cell lines, adult primary cells, such as hematopoietic stem/progenitor cells (HSPCs) and CAR-T cells, and even in embryos^8, 12–14^.

p53 inhibitor^15^ and DNA-PKcs inhibitors (AZD7648)^16^ are being explored to improve homology-direct repair (HDR) efficiency. However, these DNA repair pathway modulations could exacerbate post-editing ON-target genotoxicity^6, 17, 18^.

In this context, the clinical deployment of CRISPR-based therapies raises significant safety concerns that must be deciphered and addressed. The first published long-term data are encouraging. Indeed, CRISPR-induced aneuploidies in CAR-T cells seem to be mitigated by long-term culture *in vitro*^8^ and after graft in patients^12, 19^. In HSPCs, kilobase-scale genotoxicity following CRISPR targeting of *BCL11A* erythroid enhancers on chromosome 2 (Chr2) appears to diminish after 16-weeks post-graft in mice^20^. However, whether extra-large (megabase-scale) ON-target genotoxicity persists long term *in vivo* and can be mitigated remains unknown.

Addressing these questions requires highly sensitive methods with single-cell resolution. Microfluidic single-cell-DNA sequencing (scDNA-seq) should be appropriate for analyzing the genome integrity but its accuracy was challenged for a long time by the presence of only two genomic copies per cell. Hundreds of genomic *loci* can now be interrogated across thousands of cells. It is used to monitor intra-tumoral heterogeneity, clonal evolution and residual disease in cancer^21, 22^. More recently, scDNA-seq revealed aneuploidies in human tumors^23^.

Here, we propose a multi-assay approach to longitudinally evaluate ON-target genotoxicity induced by CRISPR-Cas9 in clinically relevant cells. We combined three orthogonal single-cell methods: (i) micronuclei (MN) detection, (ii) detection of LOH using our in-house Fluorescence-Assisted Megabase-scale Rearrangements Detection (FAMReD)^17^ assay and (iii) scDNA-seq. Importantly, by gathering SNP (single nucleotide polymorphism) and copy number variations (CNV) analyses, we demonstrate for the first time that scSNP-DNAseq is a powerful assay to measure chromosomal instability following CRISPR editing, in particular CN-LOHs with high sensitivity, *in vitro* and *in vivo*.

Our orthogonal approach offers a sensitive and robust platform for assessing genome integrity of CRISPR-edited cells and ON-target genotoxicity. It enabled us to precisely detect chromosomal instability, to map outcomes in human primary cells *in vitro*, to monitor its persistence *in vivo* after engraftment and to evaluate mitigation strategies.

## RESULTS

### Design of the scSNP-DNAseq library and method validation for LOH quantification

To detect ON-target genotoxicity induced by CRISPR-Cas9, specifically kilobase- and megabase-scale LOH, either due to a deletion (copy-loss, CL-LOH) or without loss of genomic material (copy-neutral, CN-LOH), we designed a custom scDNA-seq library, focusing on single nucleotide polymorphisms (SNPs) in Chr10/Chr11 (Fig. 1a). We first identified SNPs common in the general population using the dbSNP database (NCBI, 150,000 SNPs) and filtered for those with global allele frequency more than 0.5 (15,000 SNPs), as well as those with high frequency in European and African populations (403 SNPs distributed along chr10/11, Fig. 1b). We then designed probes to obtain 403 amplicons per cell containing the identified SNPs, with higher concentration near the CRISPR cut sites (Fig. 1b). We validated selected amplicons *in silico* by Bcftools on GIAB Coriell. Amplicons on Chr10 served as control for CRISPR targeting of Chr11, and vice-versa. After CRISPR-Cas9-editing, each amplicon from each cell was sequenced to assess genome integrity. Loss of SNPs (genotyped in unedited control cells) was used to identify and map LOH presence in each cell.

**Fig. 1:**
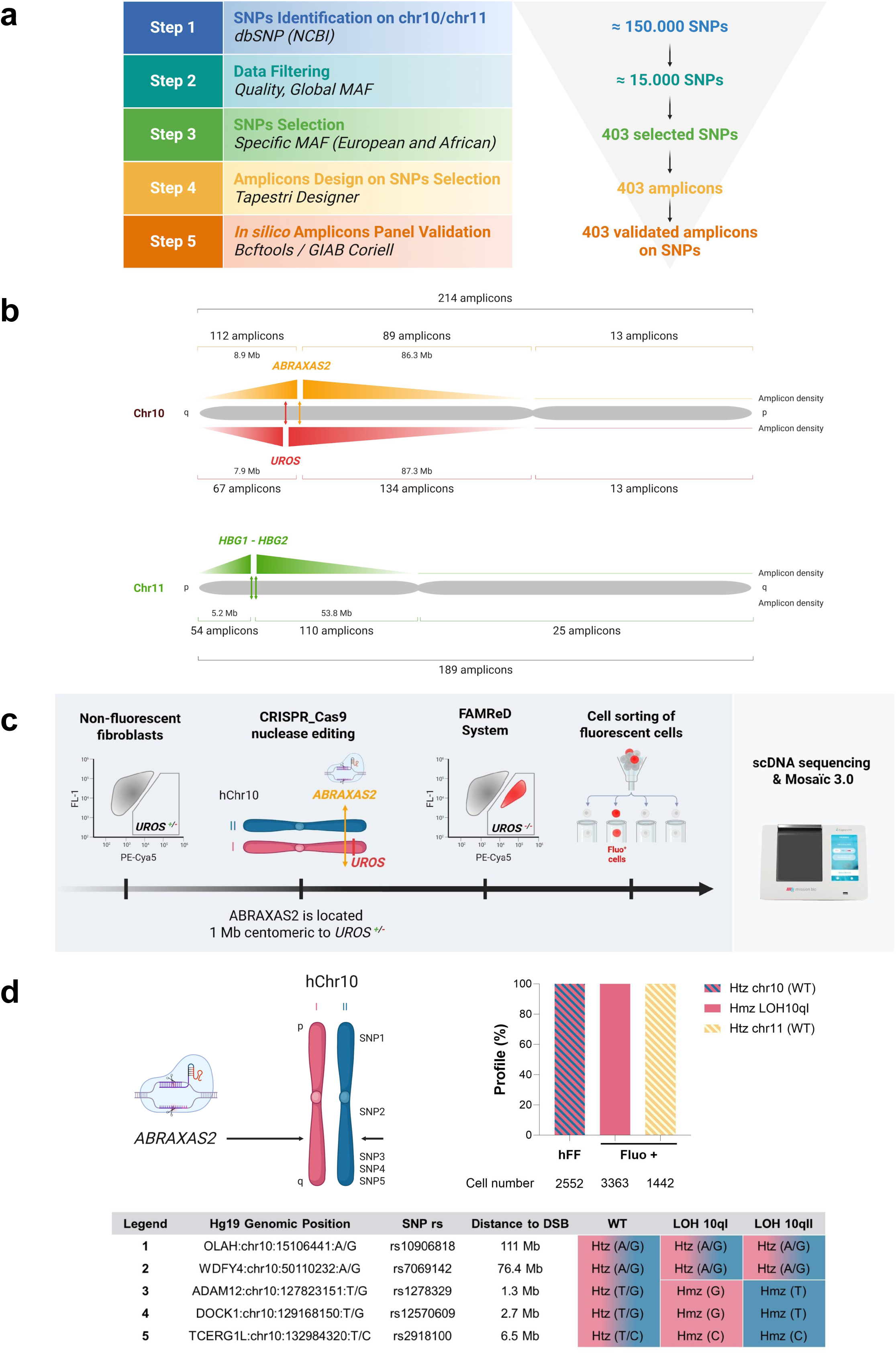
scSNP-DNAseq design to assay CRISPR-induced ON-target genotoxicity in chromosomes 10 and 11 (Chr10/11), and method validation for LOH quantification. **a,** Design of home-made scSNP-DNAseq library in Chr10/11 focusing on frequent SNPs. b, Mapping of Chr10/11 by amplicon density. Most amplicons are in the targeted chromosome arm, close to the breaking point (*ABRAXAS*2, *UROS* and *HBG1/2p*). c, Experimental design of method validation using hFAMRed system (based on *UROS*^+/-^ heterozygous hFFs). *UROS* ^+/-^ hFFs switched from non-fluorescent to fluorescent, in the event of CRISPR-induced Chr10q telomeric LOH encompassing *UROS*. *ABRAXAS2* targeting by CRISPR-Cas9 in *UROS* ^+/-^ hFFs induced fluorescent cells 15 days after editing (0.1 to 0.2%). Fluorescent cells were sorted and cultured for 2 months to obtain enough cells before analysis by scSNP-DNAseq and Mosaic 3 software. d, Top Left panel, Chr10 representation with *ABRAXAS2*, located 1 Mb centromeric to *UROS* and the five SNPs used to detect LOHs by scDNA-seq. Top Right panel, histogram of SNP profile Chr10/11 in unedited hFFs and edited fluorescent hFFs (fluo +). scSNP-DNAseq confirmed the disappearance of SNP 3/4/5 and the presence of only 1 LOH profile (LOH10qI with homozygous terminal profile) in fluorescent cells (fluo +, pink) in chr10 without losses of heterozygosity in chr11 (control chromosome in yellow). Lower panel, different SNP profiles identified by sc-SNP-DNA seq (WT, LOH 10qI, LOH 10qII) with SNP genomic position, SNP rs (reference SNP cluster ID), and distance to the DSB are indicated.

For method validation, we created a positive control cell line with a 5Mb-terminal LOH on Chr10q (LOH^+^). This was achieved using our previously published FAMReD system^17^, based on human heterozygous *UROS*^+/-^ fibroblasts (hFFs) (Fig. 1c). By targeting *ABRAXAS2* by CRISPR nuclease, located 1Mb centromeric to the *UROS* locus in Chr10q, we observed a slight increase in the proportion of fluorescent cells 15 days post-editing (0.1 to 0.2%), suggesting the occurrence of rare LOHs events at *UROS locus*, leading to the loss of the functional *UROS^+^* allele, and subsequent porphyrins accumulation. We then sorted the fluorescent cells (Fig. 1c), and analyzed them using scSNP-DNAseq with Mosaic 3.0 software. We identified SNPs located along Chr10q and selected five well-sequenced SNPs in *UROS*^+/-^ hFFs before editing (Fig. 1d). As expected, more than 99% of cells exhibited an extra-large LOH extending from the CRISPR cut-site (*ABRAXAS2*) to the telomere. We identified a unique LOH event, characterized by an allele carrying the *UROS* mutation (LOH10qI), thereby validating the method at the allele level (Fig. 1d). Importantly, the non-targeted Chr11 remained fully heterozygous, with no SNP loss.

### scSNP-DNAseq and global UMAP reveals frequent ON-target genotoxicity after double-cut in *HBG1/2* promoters in HSPCs

To investigate CRISPR-induced ON-target genotoxicity in human HSPCs, we edited the cells with the gRNA-68, which targets the two *HBG1/2* promoters, as previously reported by Sharma et al^24^. This protocol is known to predominantly induce 5kB-deletion between the two CRISPR-induced cut sites. We confirmed editing efficiency by long-range PCR and NGS analysis (Nanopore sequencing, Supplementary Fig. 1a). In parallel, we performed scSNP-DNAseq and evaluated genome rearrangements using Uniform Manifold Approximation and Projection (UMAP) for dimensionality reduction and cluster identification (Fig. 2a). First, we established the initial SNP profile in the unedited HSPCs. Using principal component analysis (PCA) and UMAP, we clustered 5161 edited cells, revealing three distinct populations based on 8 components and 88 detected SNPs (Fig. 2b). The majority of the cells (orange in UMAP) were heterozygous for all sequenced SNPs (HET = A/B, white in the heatmap, right panel). UMAP analysis revealed two equally sized minor populations (2.4% and 2.2%, red and green respectively). In these cells, 18 SNPs located between the *HBG1/2p* cut sites and the Chr11p telomere were lost and became homozygous (either A/A or B/B, indicated by pink or blue respectively, Fig. 2b right), mapping an extra-large LOH (∼ 5Mb). These abnormal cells were classified as LOH11pI and LOH11pII, respectively.

**Fig. 2:**
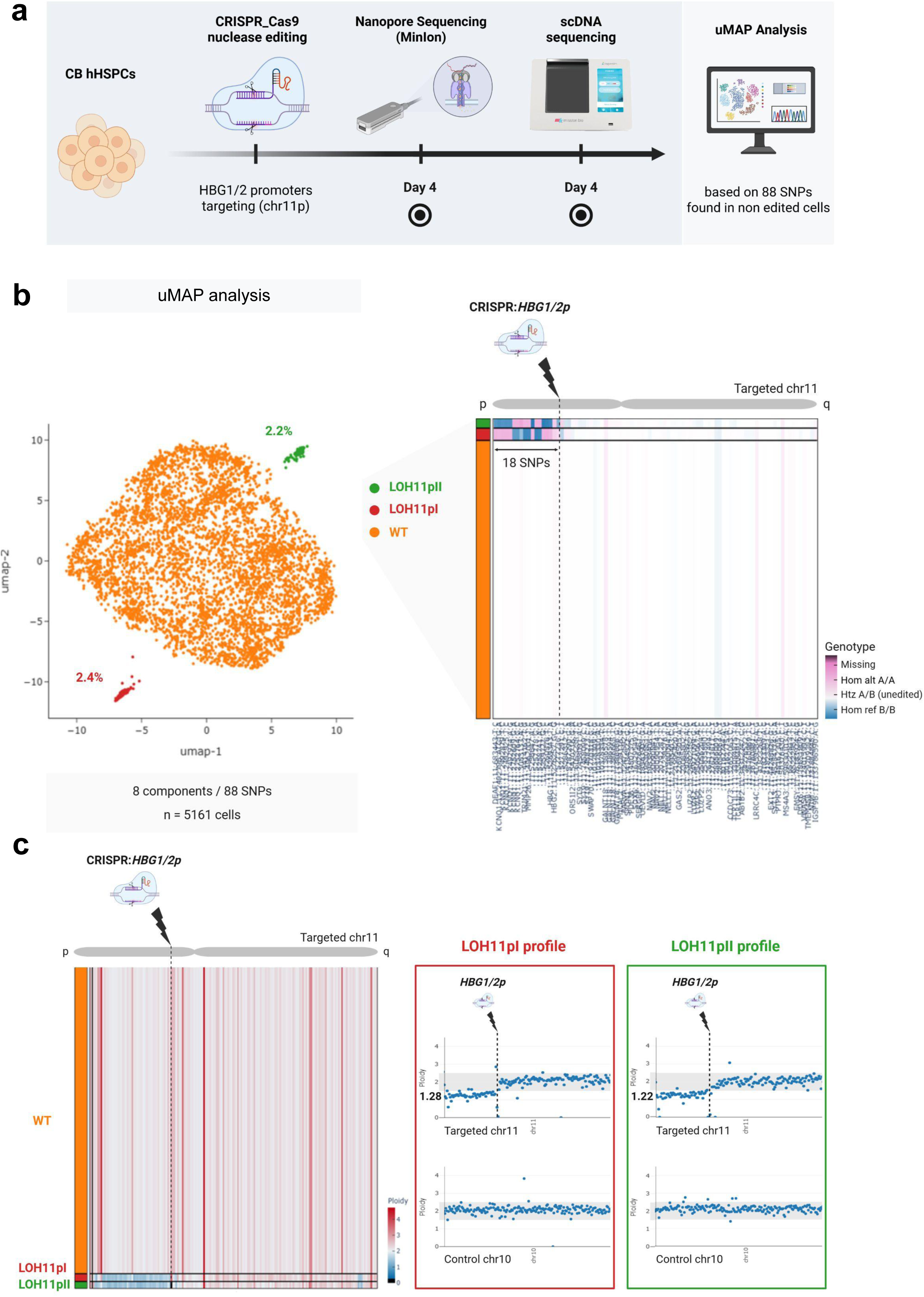
Global UMAP clustering and LOH detection following *HBG1/2* promoters targeting by scSNP-DNAseq in HSPCs. **a**, Experimental workflow. *HBG1/2* promoters were targeted using CRISPR-Cas9 in human HSPCs from cord blood (CB), followed by single-cell DNA sequencing (scDNA-seq) analysis on day 4. Dimensionality reduction was performed using principal component analysis (PCA) to capture major variance components, followed by uniform manifold approximation and projection (UMAP) for non-linear embedding. Clustering was performed using hierarchical density-based spatial clustering of applications with noise (HDBSCAN) with a minimum cluster size of 22, enabling the identification of distinct genomic subpopulations. **b**, UMAP projection and SNP-based classification of single-cells. **Left panel**, UMAP representation of 5,161 analyzed cells (99.47% retained), with three distinct profiles: no LOH (orange), LOH11pI (red), and LOH11pII (green). **Right panel**, Convolved heatmap displaying all heterozygous SNPs in cell population on chr11 diversity across groups. Orange cells from UMAP are heterozygous (white in the heatmap A/B) for all analyzed SNPs whereas green and red cells from UMAP have lost heterozygosity (18 SNPs lost, pink (Hom A/A) or blue (Hom B/B) in the heatmap between the *HBG1/2p* cut sites and the telomere). **c,** Ploidy analysis of LOH events. Averages of read depth of each amplicon were calculated in non-rearranged cells (“WT”) and used as a reference for a ploidy of 2. **Left panel**, heatmap representation of CNV across samples, highlighting copy-loss LOH regions (blue). **Right panel**, cGH-like chromosomal copy number profiles for LOH11pI (red box) and LOH11pII (green box), showing a reduction in ploidy (respectively 1.28 and 1.22, mean) in the telomeric part of targeted Chr11p arm. The position of the DSB is indicated. The non-targeted Chr10 was used as a control. Ploidy in LOH genomic area was calculated by the mean ratio of read depth of amplicons in the LOH area of edited LOH cells to non-edited cells.

To classify these LOHs as either copy-neutral (CN-LOH) or copy-loss (CL-LOH), we analyzed amplicon read-depth using the Mission Bio Mosaic pipeline to infer copy number variations (CNVs). Ploidy was calculated for each amplicon and cell exhibiting LOH by normalizing read counts against the mean of read counts of the “normal/orange” cell population. We observed abnormal ploidy at the chr11p extremity in both LOH populations (LOH11pI and LOH11pII, Fig. 2c left). Notably, genomic area with reduced ploidy (indicated in blue in the heatmap) corresponded to extra-large LOH events spanning from the cut site to the telomere. LOH ploidy was estimated by averaging the ploidy of amplicons within the LOH genomic region for each cell in both LOH populations (red and green). Ploidy in the LOH area was reduced: 1.28 and 1.22 for LOH11pI and LOH11pII respectively (Fig. 2c right) compared to a ploidy of around 2 (normal) in the centromeric region of 11p and 11q arm, and the control chr10. Copy-loss LOHs would be expected to show a ploidy of 1. These results suggest a mixture of CN-LOH and CL-LOH events. In summary, UMAP-based SNP analysis revealed significant ON-target genotoxicity following *HBG1/2*p targeting by CRISPR, with 4.6% of HSPCs exhibiting megabase-scale terminal LOH on Chr11p, the majority of which were terminal deletions (∼ 70%).

### scSNP-DNAseq focused on high-quality SNP confirms terminal copy-loss LOH and reveals interstitial LOH in HSPCs

To further explore in-depth megabase-scale LOHs (5 Mb), we reanalyzed the same edited cells, focusing on five high-quality SNPs along the targeted chr11 (selected from the 88 SNPs previously identified), each with a high genotype call rate (>60% of cells) (SNP1-5, megabasic panel, Fig. 3a/c). Using this refined method, SNP profile clustering revealed a substantial percentage of cells (7.4%) exhibiting loss of the SNP 3-5, located between the *HBG1/2*p cut sites and the 11p telomere. This finding confirms the presence of the two distinct telomeric LOHs (LOH11pI and LOH11pII) in HSPCs 4 days after transfection (D4, Fig 3b/c). This method corroborates the global genotoxicity patterns identified *via* UMAP-based clustering. Yet it demonstrates increased sensitivity, and highlights a higher frequency of chromosomal rearrangements (7.4 vs 4.6%).

**Fig. 3:**
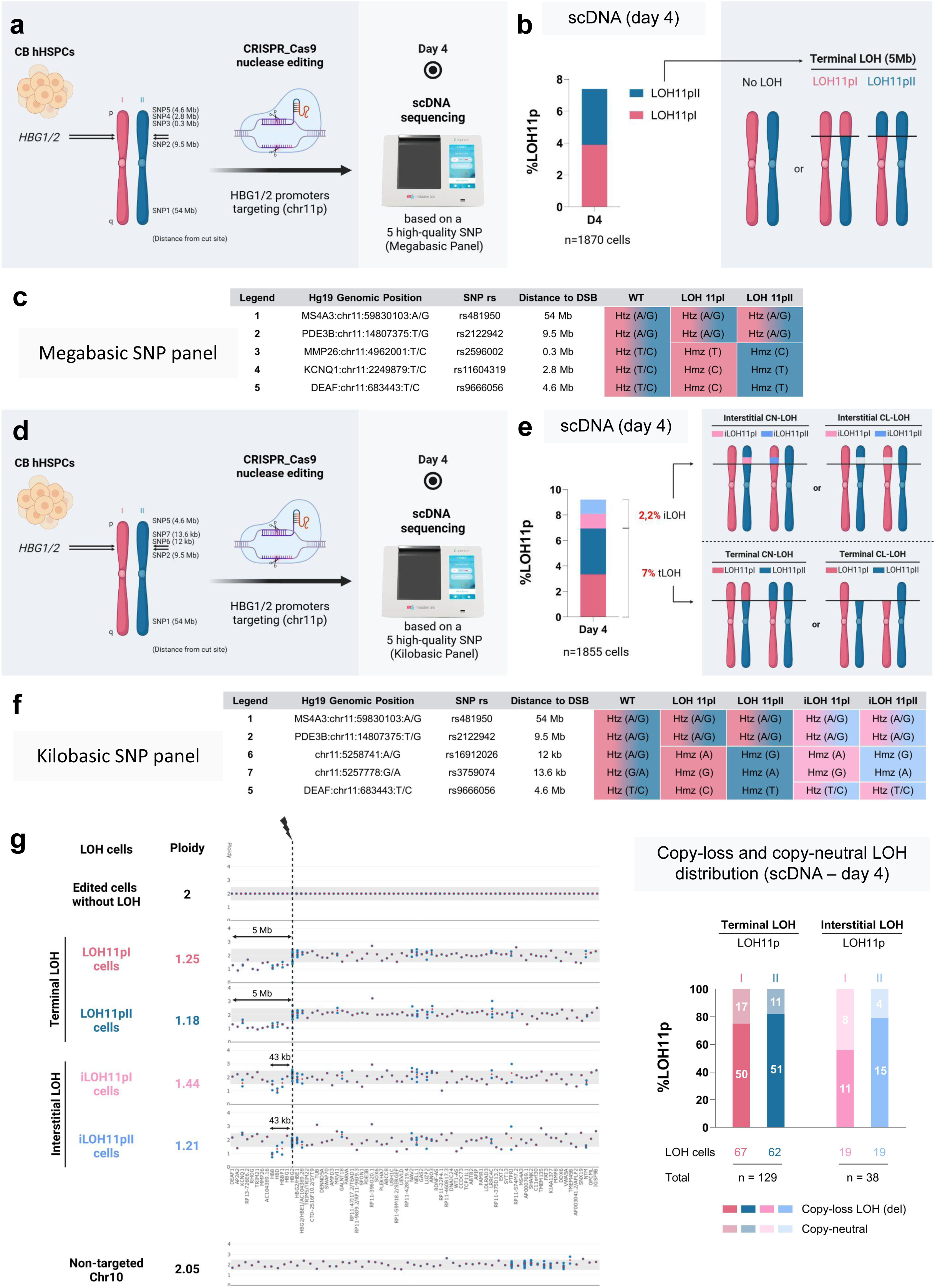
Genotoxicity analysis using high-quality SNP-amplicons revealed that *HBG1/2* promoter targeting by CRISPR induces a wide range of LOHs in HSPCs. **a,** experimental design. HSPCs cells from cord blood (CB) were transfected with Cas9 nuclease RNP and gRNA targeting *HBG1/2* promoters and scSNP-DNAseq was performed at day 4. Five high-quality SNPs along the Chr11 were chosen (megabasic SNP panel), one on the non-targeted region 11q (SNP1), one in Chr11p between the centromere and the cut site *HBG1/2p* (SNP2) and three telomeric to the cut site at 0.3, 2.8 and 4.6 Mb, (SNP 3-5). **b,** Histogram represents the % of LOH at day 4 after transfection and revealed the presence of 3 LOH profiles (WT = no LOH, telomeric LOH11pI and LOH11pII with loss of SNP3-5 (red or blue allele respectively). **c,** different SNP profiles identified by sc-SNP-DNA seq and megabasic SNP panel pipeline (WT, LOH 11pI, LOH 11pII) with SNP genomic position, SNP rs, and distance to the DSB are indicated. **d,** a novel set of SNPs was chosen using two interstitial SNPs (6 and 7) on the telomeric side (kilobasic SNP panel). **e**, scSNP-DNAseq and kilobasic SNP panel pipeline revealed the presence of 2 novel interstitial LOH profiles (iLOH). Histogram represents the % of interstitial and telomeric LOH (iLOH and tLOH) at day 4 after transfection. **f,** different SNP profiles identified by sc-SNP-DNA seq and kilobasic SNP panel pipeline (WT = no LOH), LOH 11pI, LOH 11pII, iLOH 11pI, iLOH 11pII) with SNP genomic position, SNP rs, and distance to the DSB are indicated. **g, left panel**, Ploidies and cGH-like representation (CNV analysis) along the targeted Chr11 for the 5 profiles (WT, LOH11pI, LOH 11pII, iLOH11pI and iLOH11pII), and of the non-targeted Chr10 (negative control). Ploidy in LOH genomic area was calculated by the mean ratio of read depth of amplicons in the LOH area of edited LOH cells to non-edited cells. **right panel,** cumulative histogram of copy-neutral and copy-loss LOHs in each profile (number of cells indicated).

Thanks to this increased resolution, we hypothesized that smaller, kilobase-scale LOHs may be present but undetected by UMAP analysis. We then looked for smaller interstitial LOHs in the same edited cells. To test this, we substitute SNP3 and SNP4, located 0.3 and 2.8Mb from the CRISPR cut sites) with SNP6 and SNP7, positioned closer to the cut sites, (12 and 13.6 kb respectively (Fig. 3d/f, kilobasic panel). Clustering based on this kilobase-resolution panel identified, in addition to previously characterized terminal LOH (n=129), two additional cell populations with new SNP profiles. Specifically, 2.2% of cells (n= 38) exhibited loss of SNP6-7 while retaining distal telomeric SNP5 (Fig. 3e/f), indicative of interstitial LOHs (iLOH11pI and iLOH11pII), equally distributed between maternal and paternal alleles. These interstitial events raised the total LOH frequency to approximately 9%. No LOH were detected on the non-targeted Chr10, supporting the specificity of CRISPR-Cas9 induced genotoxicity. To determine whether the observed LOH were due to copy number loss (CL) or copy-neutral (CN) events, we assessed CNVs in LOH-containing cells using the kilobasic SNP panel (Fig. 3g), considering all amplicons. LOH ploidy was estimated by the average of amplicon ploidies across the LOH region for each cell in each population. Among the 1855 analyzed cells, 69 cells exhibit a terminal 11p LOH. In these cells (LOH11pI and LOH11pII), the calculated mean LOH ploidies were 1.25 and 1.18 respectively, suggesting that ∼ 80% of these LOHs resulted from terminal deletions (Fig. 3g, day 4). In the 38 cells with interstitial LOHs (iLOH11pI and iLOH11pII), mean ploidies in the affected 43kb LOH area (from the cut site to *AC104389.16*), were 1.44 and 1.21, respectively, again indicating a predominance of interstitial deletions (Fig. 3g). Altogether, we demonstrated the value of high quality SNP-focused scDNA-seq for deciphering and precisely mapping the ON-target genotoxicity of *HBG1/2p* targeting by CRISPR-Cas9 in primary cells. Our data revealed a high frequency of both interstitial and terminal deletions at day 4.

### AZD7648 increases short-term genotoxicity in human fibroblasts (hFFs)

Pharmacological modulators known as “HDR boosters” have been proposed to improve the HDR rate during CRISPR-mediated genome editing. To evaluate the impact of three HDR boosters on editing efficiency and genotoxicity, we treated hFFs with XL413, an inhibitor of CDC7 and two DNA-PKcs inhibitors, NU7441 and AZD7648, during *ABRAXAS2* editing with a high-fidelity (HiFi) Cas9, and a single-strand oligo-desoxy ribonucleotide (ssODN) template to obtain HDR (Fig. 4a). We monitored ON-target genotoxicity by a multiplex single-cell method approach combining micronuclei assays, cytometry-based LOH detection and scSNP-DNAseq.

**Fig. 4:**
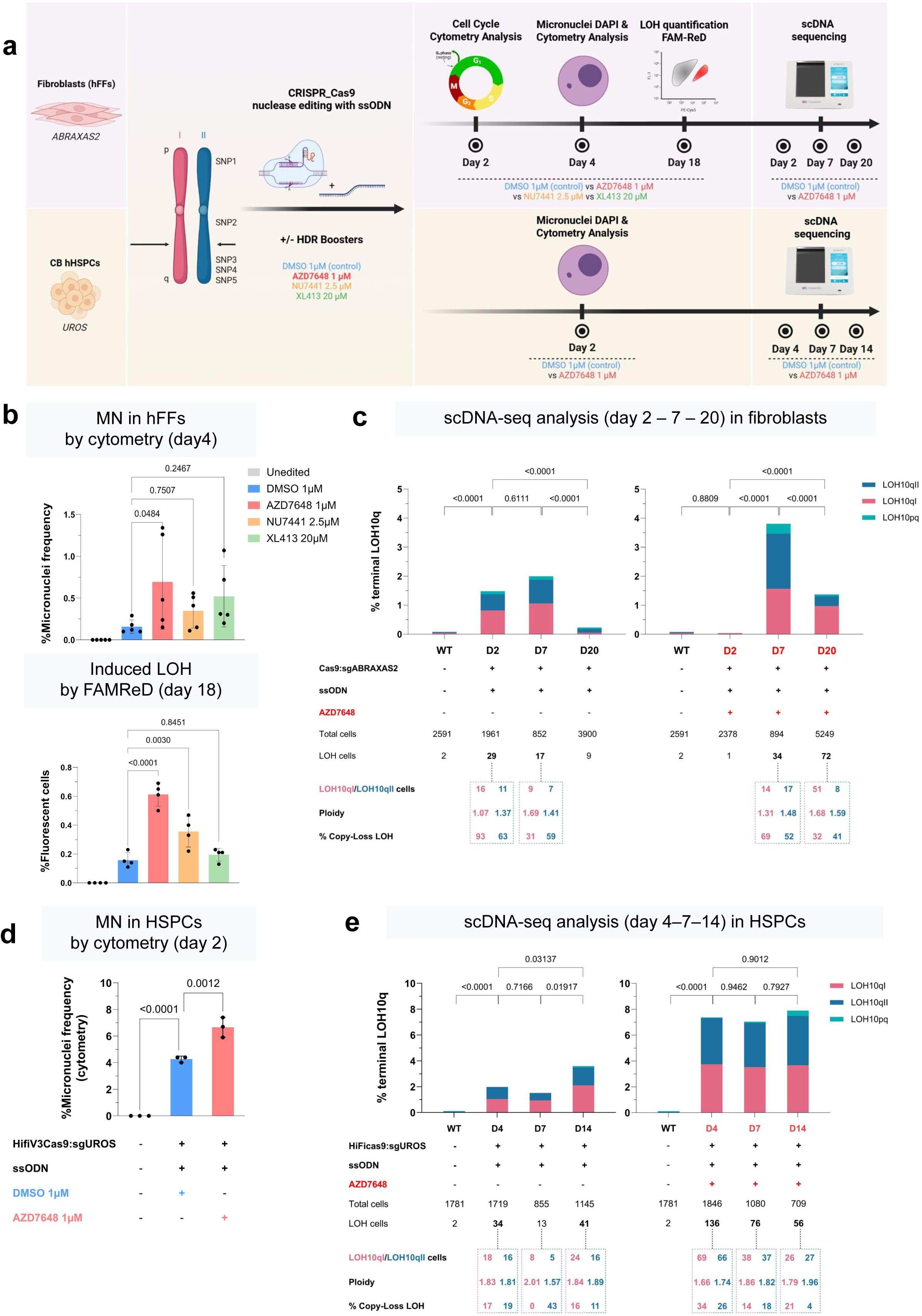
Time course of AZD7648 short-term megabase-scale genotoxicity in hFFs ans HSPCs. **a,** experimental design. hFFs and HSPCs were transfected with Cas9 RNP, ssODN template and gRNA targeting *ABRAXAS2 or UROS,* with or without HDR boosters 18 h before transfection and for 2 days after transfection (AZD7648, in red: 1 µM; NU7441, in orange: 2.5µM; XL413, in green: 20µM; DMSO control in blue). Schema of the Chr10 and analyzed SNPs. After transfection, cell cycle phases (day 2), micronuclei count (day 4), editing efficacy (day 7), LOH detection by FAMReD (day 18, only for hFFs) and scSNP-DNAseq kinetic were performed. **b, top panel,** % of micronuclei (MN) quantified by cytometry at day 4 (Mean ± SD, n=5 independent experiments, One-way ANOVA test, at least 10,000 cells per condition) in hFFs edited at the *ABRAXAS2 locus*. DMSO was used as a control of the 3 HDR boosters. **Bottom pane**l, % of fluorescent cells induced by *ABRAXAS2* editing at day 18 using the FAMReD system. Fluorescent cells are due to telomeric megabase-scale LOH (Mean ± SD, n=4 independent experiments, One-way ANOVA test). **c**, 5 high-quality SNPs along the Chr10, present in unedited cells, were selected. % of LOH in HFFs detected by scSNP-DNAseq at day 2, 7 and 20 after transfection, with or without CRISPR-Cas9 and AZD7648 exposure (right histogram with AZD7648). Percentages of LOHs in hFFs were compared by the Chi-square test. Number of cells exhibiting LOH10pI and LOH10pII profiles, ploidy and % of copy-loss LOH are indicated. **d**, histogram with % of MN quantified by cytometry at day 2 after HSPCs editing at the *UROS* locus (Mean ± SD, at least 10,000 cells per condition, n=3 independent experiments, One-way ANOVA test). **e**, 5 high-quality SNPs along the Chr10, identified in unedited cells, were selected. % of LOH in HSPCs detected by scSNP-DNAseq at day 4, 7 and 14 after transfection, with or without CRISPR-Cas9 and AZD7648 exposure (right histogram with AZD7648). Percentages of LOHs in HSPCs were compared by the Chi-square test. Number of cells exhibiting LOH10pI and LOH10pII profiles, ploidy and % of copy-loss LOH are indicated.

PCR and Sanger sequencing revealed that, among these tested compounds, only AZD7648 significantly increases the HDR/NHEJ ratio in hFFs (supplementary Fig. S2a/b). The “HDR-boosters” genotoxicity was first assessed using a cytometry-based MN assay at day 4 post-editing (Fig. 4b top panel). AZD7648 treatment markedly elevated MN formation after *ABRAXAS2* editing (up to 0.7% of MN/nucleus *vs* 0.16% in edited hFFs without AZD7648, fig. 4b), indicating increased genotoxic stress.

To confirm this result, we edited hFFs at *HBG1/2p* loci. Once again, an increase of micronuclei was observed in the presence of AZD7648 (supplementary Fig. 2d), without any change in editing efficiency as measured by nanopore sequencing (supplementary Fig. 2c). The FAMReD system, developed in our lab to sensitively detect telomeric LOH by cytometry following CRISPR-Cas9 editing, confirmed the genotoxicity of AZD7648 in hFFs and showed persistence of megabase-scale terminal LOH at day 18 (0.6% of fluorescent LOH^+^ cells vs 0.16% in DMSO control) (Fig. 4 bottom panel). Since FAMReD detects only 50% of the events, the true long-term LOH frequency is double (1.2% of edited cells treated with AZD7648). A smaller genotoxic effect was observed with Nu7441 (by both MN and FAMReD analyses). The effect of AZD7648 was exacerbated in *TP53*^-/-^ hFFs (supplementary Fig. 2e left, 6% of cells with telomeric LOH).

To quantify, map, and characterize the nature of post-editing LOH events in the presence or absence of AZD7648, the most genotoxic HDR booster, we performed a kinetic analysis of ON-target genotoxicity using scSNP-DNAseq analysis, before and after hFF editing at *ABRAXAS2 locus*, with a ssODN template, with or without AZD7648 (Fig. 4a, Days 2-7-20). Bioinformatic analysis first identified five informative SNPs in non edited cells, selected with stringent quality criteria to enable megabase-scale resolution, as described in Fig. 3. After editing, clustering based on these five SNPs revealed four cell populations with distinct SNP profiles (supplementary Fig. 3a): one population with heterozygous SNPs along the Chr10 (“WT”), two abnormal populations with allele loss for 3 SNPs 3/4/5, from each parental chromosome (designated telomeric “LOH10qI” and “LOH10qII”), and a fourth abnormal population showing loss of all analyzed SNP (Chr10pq SNP loss).

After editing in hFFs without AZD7648, we observed, at day 2, 1.5% of cells with telomeric 10q LOH, extending from the cut site to the telomere (Fig. 4c left upper panel D2), and rare cells with complete loss of SNPs. CNVs analysis, performed only when more than five cells were available per population, revealed reduced ploidy in both LOH11pI and LOH11pII populations (1.07 and 1.37 in LOH10qI and LOH10qII respectively), demonstrating that most LOH events were due to genomic material loss (93% and 63% of deletions (CL-LOH), in LOH10qI and LOH10qII respectively, Fig. 4c). The high rate of telomeric LOH persisted at day 7 (in 1.9% of cells) (Fig. 4c left panel), with a balanced mix of CN-LOH and CL-LOH (ploidy 1.69 et 1.41). Interestingly, in the same edited cells, the frequency of LOH dropped clearly by day 20 (0.23%).

Unexpectedly, in the presence of AZD7648, no LOH was detected on day 2 using scSNP-DNAseq (Fig. 4c right panel). We hypothesized that DNAPKcs inhibition during a DSB induction might slow down cell division, delaying the onset of ON-target genotoxicity. We therefore analyzed the cell cycle of hFFs exposed to AZD7648. When a DSB occurs concomitantly with AZD7648 treatment, hFFs tend to accumulate in the G0 phase at day 2 (Supplementary Fig. 3b). These results may explain the absence of detectable LOH immediately after editing. At day 7, however, a high rate of LOH was observed (3.9% of cells), with a mixture of CN-LOH and CL-LOH (ploidy 1.31 et 1.48). This genotoxicity was partially maintained at day 20 (1.37%), suggesting a delayed onset and sustained maintenance, compared to without AZD7648. At day 20, in the presence of AZD7648, the persistent LOHs were predominantly copy-neutral (ploidy 1.68 and 1.59, Fig. 4c right panel). “cGH like” schematic representations are provided in Supplementary Fig. 4. No LOH was detected on the non-targeted Chr11, even with AZD7648 (not shown).

Among all tested HDR enhancers, AZD7648 reached the highest short-term genotoxicity in hFFs. Kinetic reveals that LOHs, initially resulting from DSB and chromosomal truncations, tend to convert into copy-neutral LOHs and diminish in frequency at day 20.

### AZD7648 increases short-term ON-target genotoxicity in HSPCs targeting *UROS* (Chr10q)

Because DNA repair pathways are cell-type dependent, we next investigated the genotoxicity of HDR boosters in HSPCs (Fig. 4a). We targeted the *UROS* locus in Chr10q in the presence or absence of HDR boosters. Nanopore NGS sequencing confirmed that AZD7648 is the most potent HDR booster, significantly increasing the HDR rate (77% *vs* 12%, supplementary Fig. 5a). Again, AZD7648 exposure during editing led to a dramatic increase in MN formation (up to 3-4% of MN/nucleus), suggesting that modulating DNA repair pathways during DSB repair also elevates genotoxicity in HSPCs (Fig. 4d and illustrative supplementary Fig 5b). DNA staining by DAPI and fluorescence microscopy of edited HPSCs further confirmed the increase in nuclear abnormalities upon AZD7648 treatment (supplementary Fig. 5c).

To monitor the kinetics of unwanted genome editing outcomes in HSPCs, we designed a time-course scSNP-DNAseq analysis with or without AZD7648. HSPCs were edited, cultured, and collected on days 4, 7 and 14 for scSNP-DNAseq. We identified four cell populations post-editing: “WT” without LOH (all SNP heterozygous), telomeric LOHs “LOH 10qI” and “LOH 10qII” (SNPs 3/4/5 loss), and whole-chromosome LOH “LOH 10pq” (all SNPs lost) (supplementary Fig. 5d). Without AZD7648, CRISPR editing led to 2-4% of rearranged cells, mostly showing telomeric LOHs (“LOH 10qI and LOH 10qII”, fig. 4e left) from day 4 to day 14, confirming that even a single DSB can induce ON-target genotoxicity. AZD7648 exposure significantly increased the proportion of cells with telomeric LOHs throughout the time-course (day 4 to day 14), reaching ∼7%, a 3-fold increase compared to CRISPR alone, fig. 4e right). These results indicate that telomeric LOHs are stable *in vitro* over a two-week period. No LOH was detected on the non-targeted Chr11, even with AZD7648. Under all conditions, with or without AZD7648, and whatever the time point, the majority of LOHs were copy-neutral, with ploidy values around 2 (ranging from 1.74 to 2, Fig. 4e). The whole-chromosome ploidy patterns “cGH-like” are shown in Supplementary Fig. 6.

Taken together, these data demonstrate that AZD7648 induces both short- and medium-term ON-target genotoxicity in HSPCs, with immediate cellular stress (micronuclei and nuclear abnormality formations) and the persistence of extra-large copy-neutral LOH events *in vitro* for up to two weeks.

### Palbociclib prevents post-editing ON-target genotoxicity in HSPCs

Mitigating ON-target genotoxicity is essential for the safe development of CRISPR-based gene therapy using nucleases. DSB-induced large-scale genotoxicity is thought to mainly occur in dividing cells. We hypothesized that G0/G1 cell cycle arrest could mitigate chromosomal rearrangements observed in HSPCs (Fig. 2-3-4d/e). In a previous work, we demonstrated that palbociclib, a CDK4/6 inhibitor, effectively blocks fibroblasts in the G0/G1 phases^17^. We therefore investigated whether palbociclib could transiently arrest HSPCs in G0/G1 during editing and thereby prevent genotoxicity in clinically relevant human HSPCs.

We first monitored its impact on (i) HSPCs cell cycle progression, (ii) editing efficiency (*via* long-range PCR and nanopore sequencing) and (iii) genotoxicity using multiplexing single-cell approaches (micronuclei and scSNP-DNAseq) (Experimental design, Fig. 5a). We optimized the dose and timing of palbociclib to transiently block HSPCs in the G0/G1 phases (Fig 5b and illustrative supplementary Fig 9a). Just after thawing, nearly all HSPCs were in G0/G1 phases (99%). While this proportion decreased after 12hours in culture, palbociclib maintained a higher fraction of cells in G0/G1 (84% vs 63% without palbociclib). A 36h-palbociclib exposure to palbociclib further enhanced this effect (92% of cells in G0/G1 phase vs 70% without palbociclib), which was reversed upon drug removal (81% G0/G1 cells with palbociclib after 36h-treatment and a 48h-clearance/washout vs. 76% in untreated cells, Fig. 5b). For comparison, cytokine starvation was also tested to reduce cell proliferation (supplementary Fig. 9b/c/d). Comparison between palbociclib *vs* cytokine starvation revealed that palbociclib was more efficient than 3 ng/mL cytokine starvation to synchronize the cells (Fig. S9b 36h). Complete cytokine starvation (0 ng/mL) is equally potent to palbociclib to arrest the cells in G0/G1 (Fig. S9b 36h) and block proliferation (Fig. S9c) but more toxic (Fig. S9d). Thus, palbociclib was selected for transient HSPCs cell cycle arrest in the remainder of the study.

**Fig. 5:**
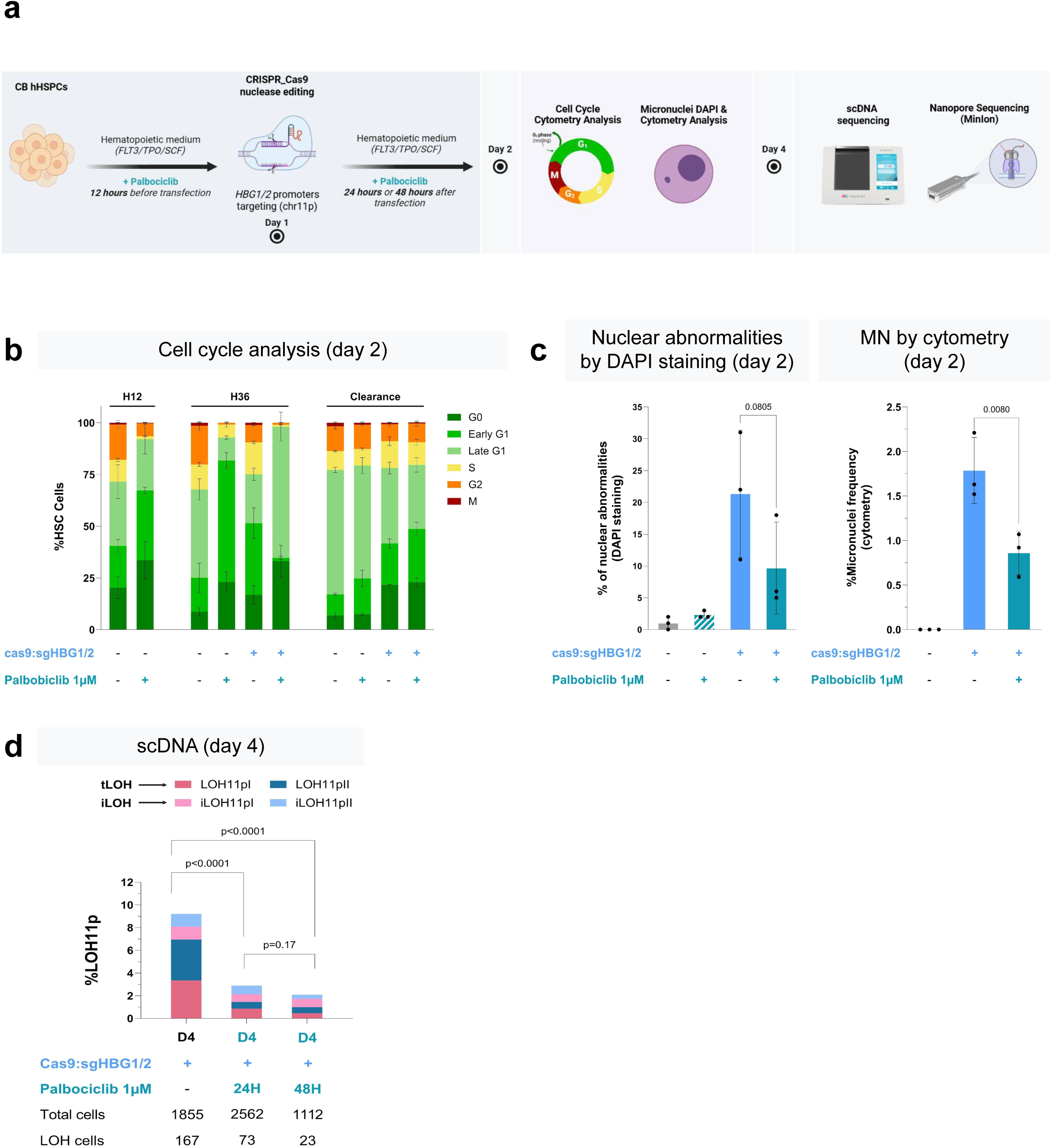
Palbociclib prevents genotoxicity in HSPCs. **a,** experimental design. HSPCs from CB were edited with Cas9 RNP and gRNA targeting *HBG1/2* promoters with or without palbociclib exposure (1 µM, overnight before transfection and for 24h or 48h post-transfection). Cell cycle analysis, nuclear abnormalities and micronuclei count (day 2), Nanopore sequencing and scSNP-DNAseq (day 4) were performed. **b,** Histogram of cell cycle after 12h and 36h of cell culture with or without palbociclib (Mean ± SD, n=3 independent experiments), and after 48h of clearance. **c, left panel,** fluorescence microscopy: % of nuclear abnormalities quantified by DAPI staining at day 2 (Mean ± SD, n=3 independent experiments, with 300, 100 and 100 cells respectively. Paired t-test). **right panel**, % of micronuclei (MN) quantified by cytometry at day 2 (Mean ± SD, n=3 independent experiments. One-way ANOVA test, at least 10,000 analyzed cells per condition). **d**, % of LOHs (interstitial and terminal) quantified by scSNP-DNAseq (using 5 high-quality SNPs, kilobasic panel in supplementary 3f.) at day 4 without or with palbociclib exposure. Histogram represents the % of interstitial and telomeric LOH (iLOH in pink and light blue, and tLOH in red and blue). Percentages of LOHs in HSPCs were compared by the Chi-square test.

We then edited *HBG1/2p* in HSPCs in the presence of palbociclib. Treatment during editing maintained > 95% of cells in G0/G1 phases (Fig. 5b) without compromising editing efficiency (supplementary Fig. 1).

To evaluate short-term ON-target genotoxicity, we first analyzed nuclear abnormalities in HSPCs at day 2 by DAPI staining (micronuclei, nuclear buds…). Fluorescence microscopy revealed that CRISPR targeting of *HBG1/2p* induced a high rate of abnormal nucleus, hallmarks of genomic instability (around 20% of cells with abnormal nucleus, fig 5c left, blue). Palbociclib reduced this frequency by more than two-fold (Fig. 5c left, green).

To confirm, we performed cytometric analysis of micronuclei (MN) by cytometry. *HBG1/2p* targeting by CRISPR led to a high MN rate (Fig. 5c right, blue), which was again reduced over 2-fold with palbociclib in HSPCs (from 1.8% to 0.9%, Fig. 5c right, green).

To track chromosomal rearrangements, we collected cells on day 4 to perform scSNP-DNAseq using the kilobasic panel described in Fig. 3f (Fig. 5d). Total LOH frequency (telomeric + kilobasic) dropped from 9.2% in edited controls to 2.9% or 2.1%, with 24h or 48h of palbociclib exposure, respectively. These data demonstrate for the first time that 24-hour exposure to palbociclib is sufficient to protect HSPCs from ON-target genotoxicity. In the remaining rearranged cells, all four LOH subtypes were still present, but palbociclib altered their distribution. Specifically, telomeric LOH frequency dropped from 6.95% to 1% after 48h of palbociclib treatment, a 7-fold reduction, while interstitial LOH frequency was only reduced two-fold (from 2.25% to 1.1%, Fig. 5d). Ploidy analysis of each LOH event (“cGH like” patterns, Supplementary Fig. 7) showed that the proportion of CL-LOH and CN-LOH was unchanged after 24h of palbociclib (ploidy ∼1.3, corresponding to 70% deletions in both conditions). Altogether, these findings demonstrate that transient G0/G1 arrest by palbociclib (12h pre-treatment + 24h post-editing exposure) effectively protects HSPCs from DSB-induced damages (preferentially megabase-scale LOH), without compromising editing efficiency via NHEJ.

We hypothesized that G0/G1 maintenance by palbociclib may prevent genotoxicity by promoting end joining. Because MMEJ is absent in G0/G1, we challenged NHEJ implication in palbociclib preventing effect. To test this, we used the highly sensitive FAMReD system to evaluate the effect of concomitant NHEJ inhibition by AZD7648 during palbociclib exposure (supplementary Fig. 8). Interestingly, in the presence of AZD7848 plus palbociclib, megabasic LOH rate was exacerbated (supplementary Fig. 8b). However cell cycle analysis showed that cells remained synchronized in G0/G1 (supplementary Fig. 8a). These data demonstrate that the protective effect of palbociclib relies on NHEJ activity to prevent megabase-scale chromosomal rearrangements. InDels profiles further supportive NHEJ importance in palbociclib-treated cells. CRISPR preferentially generated small InDels (<5 bp, typical of NHEJ) rather than larger InDels (> 5bp, typical of MMEJ) (supplementary Fig. 4c).

### Palbociclib improves cell graft capability and hematopoietic reconstitution of HSPCs

By inducing G0/G1 cell cycle arrest in HSPCs, palbociclib could theoretically impair cell proliferation and impact *in vitro*/*in vivo* hematopoietic function (Fig. 6a). We first carried out an in *vitro* test of proliferation. As expected, palbociclib transiently reduced HSPCs proliferation in hematopoietic liquid culture (Fig. 6b), similar to complete cytokine starvation (0 ng/mL, red line, supplementary Fig. 9c) but with less toxicity (0 ng/mL, 53.5 % of alive cells *vs* 69% using palbociclib and even 78% using palbociclib in edited cells, supplementary Fig. 9d).

**Fig 6:**
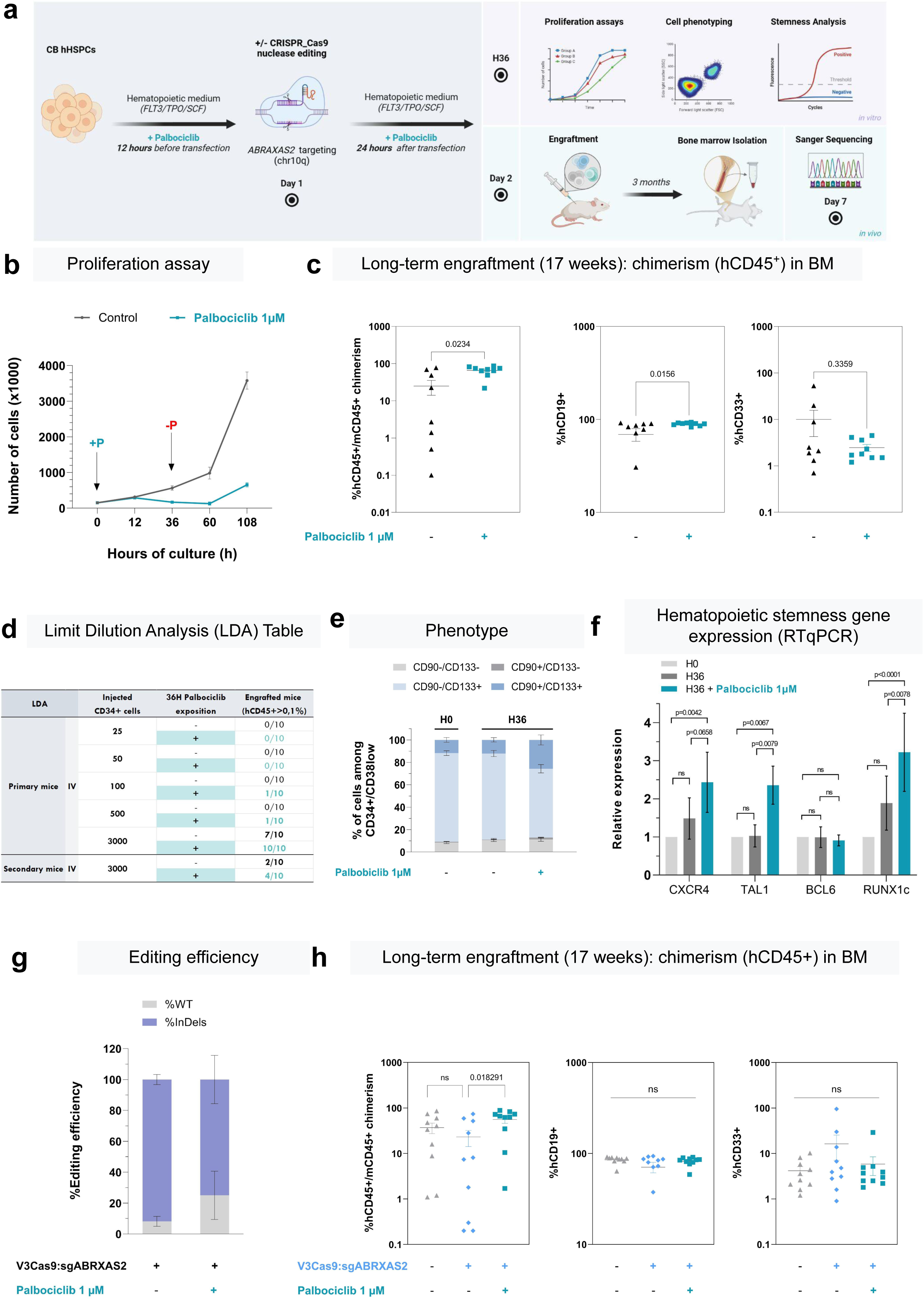
Palbociclib effect on HSPCs phenotype and graft capability. **a,** experimental design. HSPCs from CB, edited or not at *ABRAXAS2 locus,* were cultured with or without palbociclib exposure (1 µM, medium with 100ng/mL cytokines, 36h after transfection). Proliferation, phenotype (surface markers and stemness gene expression) and long-term engraftment in NSG mice were performed. **b**, Proliferation assay by cell count during 108h, with or without palbociclib added during the first 36h (Mean ± SD, n=3 independent experiments). **c,** Analysis of BM cells 17 weeks after engraftment. **left panel**, Chimerism (% of hCD45^+^ cells) in the BM of engrafted mice after injection of 10000 cells, pre-incubated with or without palbociclib (1µM, 36h) (Mean ± SEM, n = 10 mice per group, Mann-Whitney test) at week 17. **Middle and left panel**, Cell distribution in hematopoietic lineages analyzed by flow cytometry in the BM: % of hCD33 and hCD19 among hCD45^+^ cells. **d**, Limiting dilution assay in NSG mice (n= 120). A scale from 25 to 3000 cells, pre-incubated with or without palbociclib (1µM, 36h) were intraveinously (IV)-injected and engraftment was evaluated at week 17 by hCD45^+^ chimerism in femurs. BM (1 femur) of mice engrafted with 3000 cells sorted were re-injected in secondary mice. Positive engraftment was considered when hCD45^+^ was > 0.1%. **e**, Cell surface marker analysis (CD34/CD38/CD90/CD133) by cytometry to evaluate hematopoietic stem population, at thawing (H0), 36h after cell culture with or without palbociclib (Mean ± SD, n=3 independent experiments). **f**, hematopoietic stemness gene expression of *CXCR4, TAL1, BCL6* and *RUNX1c* by RTqPCR. Results are normalized on expression in HSPCs at thawing (H0) (mean ± SD, n=3 independent experiments, two-way ANOVA). **g,** Editing efficiency at day 7 evaluated by sanger sequencing and DECODR software. **h,** Analyze of hematopoietic reconstitution in NSG mice 17 weeks after engraftment of *ABRAXAS2*-edited cells, without or with palbociclib. BM from femur was analyzed (hCD45, hCD19 and hCD33) (Mean ± SEM, n=10 mice, Mann-Whitney test).

This proliferation slow-down could affect *in vivo* HSPCs properties. To assess longer-term consequences, we grafted equivalent numbers of HSPCs pre-treated or not with palbociclib into NOD-*scid* IL2Rgammanull (NSG) mice (experimental design Fig. 6a). After 17 weeks post-transplantation, bone marrow (BM) chimerism (% hCD45^+^ cells) was higher in mice receiving palbociclib-treated cells (Fig. 6c left). Hematopoietic lineage reconstitution (hCD33^+^ and hCD19^+^ cells) in BM was comparable between conditions, demonstrating that palbociclib does not impair multilineage differentiation (Fig. 6c, middle and right panels).

To quantify grafting capacity, we performed limiting dilution analysis (LDA) across 120 mice (Fig. 6d). LDA confirmed that palbociclib does not compromise HSPCs engraftment and even enhances it. Indeed, the number of transplanted mice is greater with palbociclib than without. For example, among mice transplanted with 100 and 500 cells, only those incubated with palbociclib show successful engraftment (2/20 vs 0/20 mice, Fig. 6d). This result was confirmed in secondary recipient mice, with a higher number of engrafted mice in the palbociclib group (4/10 vs 2/10, Fig. 6d). The frequency of SCID Repopulating Cells (SRCs) was 2.6-fold higher with palbociclib (1/1360, CI [1/740-1/2500]), compared to untreated controls (1/3541, CI [1/1700-1/7350]).

These findings suggest that, by forcing HSPCs to be in the G0/G1 phases, palbociclib could modulate their stemness properties. Phenotypic analysis after 36h of exposure showed an increased frequency of stem cell population (CD34^+^/CD38^low^/CD90^+^/CD133^+^) (from 12% to 26%, Fig. 6e), suggesting maintenance or selection of stem cells, or possible progenitor dedifferentiation. This is supported by the up-regulation of hematopoietic stemness-associated genes such as *CXCR4, TAL1* and *RUNX1* in HSPCs treated with palbociclib for 36h (Fig. 6f).

To confirm the palbociclib’s positive effect on engraftment in context of nuclease-based genome editing, we targeted *ABRAXAS2* locus in HSPCs, with or without palbociclib exposure (12h pre-treatment + 24h post-editing). Editing efficiencies were very high in both conditions (Fig. 6g). Following palbociclib removal, equal numbers of cells were transplanted into NSG mice. Again, chimerism was higher in the palbociclib group (Fig. 6h left), without altering CD19^+^ and CD33^+^ lineage distribution (Fig. 6h, middle and right panels). Together, these results demonstrate that palbociclib is not deleterious, and in fact enhances HSPCs engraftment. Its use during CRISPR protocols could therefore be explored as a strategy to prevent ON-target genotoxicity while preserving or enhancing edited-HSPCs transplantation.

### Chromosomal rearrangements disappear after long-term engraftment

To evaluate longer-term impact of LOH events on HSPCs proliferation and function, we engrafted CRISPR-edited HSPCs (targeting in *HBG1/2p)* into NSG mice for *in vivo* genotoxicity monitoring (Fig. 7a). Prior graft, we confirmed editing efficiency (Fig. 7c and supplementary 10) and the immediate genotoxicity by micronuclei assay (Fig. 7b).

**Fig. 7:**
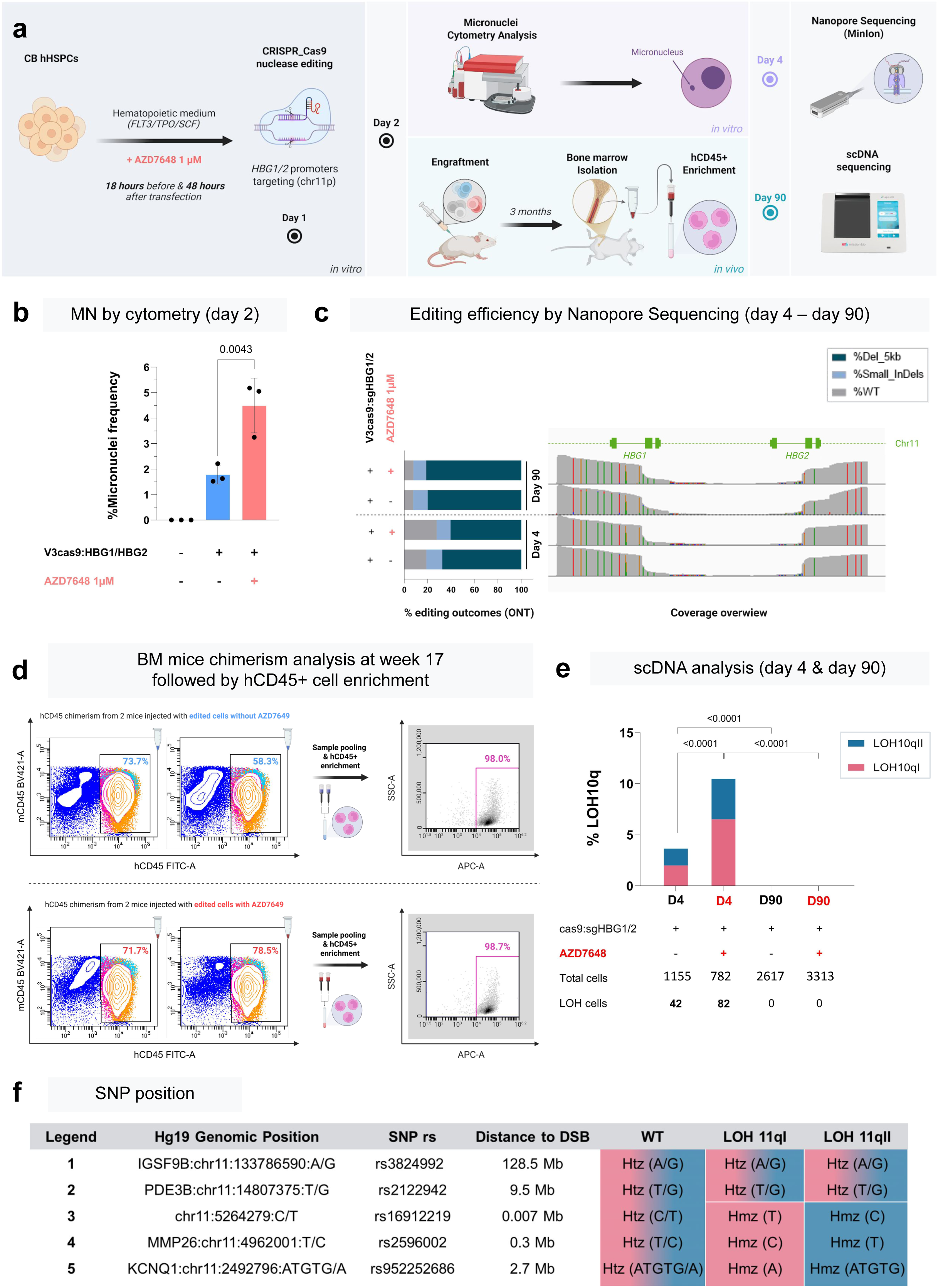
Long-term i*n vivo* genotoxicity assessment of *HBG1/2p* editing protocol, with or without AZD7648. **a,** experimental design. Human HSPCs cells from cord blood (CB) were transfected with Cas9 nuclease RNP and gRNA targeting *HBG1/2* promoters, with or without AZD7648. *In vitro* studies: micronuclei analysis (day 2), editing efficiency by Nanopore sequencing (day 4) and scSNP-DNAseq (day 4). *In vivo* study: Engraftment of 400 000 edited human HSPCs at day 2 in 4 NSG mice (2 with and 2 without AZD7648). BM harvesting, sorting of hCD45^+^ cells at day 90, editing efficiency by Nanopore sequencing and scSNP-DNAseq analysis to assess long-term *in vivo* genotoxicity (day 90). **b**, % of micronuclei (MN) quantified by cytometry at day 2 (Mean ± SD, n=3 independent experiments, One-way ANOVA test, at least 10,000 cells per condition) without (blue) or with (red) AZD7648. **c**, Long-range PCR and Nanopore sequencing of *HBG1/2* promoters from edited HSPCs. Histogram of % of WT, InDels and deleted alleles. Double DSB in *HBG1p* and *HBG2p* mostly induces a 5kB deletion. **d**, hCD45^+^ quantification by cytometry to measure BM mice chimerism analysis at week 17 and after hCD45^+^ cell enrichment. **e**, scSNP-DNAseq at D4 (*in vitro*, short-term), and D90 (*in vivo*, long-term) with or without AZD7648 during editing. % of telomeric LOH are indicated. Percentages of LOHs in HSPCs were compared by the Chi-square test.

We grafted 400,000 edited human HSPCs per mouse, 24 hours after CRISPR editing, with or without AZD7648 exposure (n=2 mice per condition). We then analyzed LOH frequency by scSNP-DNAseq in “before graft” and 3 months after graft (*in vivo*) to assess persistence of genotoxicity in bone marrow (BM) HSPCs (Fig. 7a). For that, we harvested BM cells from mice 90 days after graft. We obtained high chimerism (hCD45^+^ cells) highlighting a good hematopoietic reconstitution (Fig. 7d). Human hematopoietic CD45^+^ cells were sorted by magnetic beads and purity confirmed before scSNP-DNAseq analysis. Before graft, 3.6% and 10.5% of cells exhibited megabase-scale telomeric LOH (from *HBG* to the 11p telomere) in the absence or presence of AZD7648, respectively (Fig. 7e/f). However, at day 90 post-engraftment, no genotoxicity was detected, even though edited alleles persisted (Fig. 7c, day 90). These findings demonstrate the complete disappearance of rearranged cells over time *in vivo*, including in the more genotoxic AZD7648 condition.

To sum up, these data establish that scDNA seq can be used to monitor *in vivo* ON-target genotoxicity and show that Chr11p LOH^+^ cells do not persist *in vivo*, even after editing with AZD7648.

## DISCUSSION

Genotoxicity is a major concern for innovative therapeutic approaches involving genome editing. To address this, we developed a multiplex single-cell approach that combines innovative scSNP-DNAseq with a micronuclei and a LOH cytometry-reporter assays. By combining CNV analysis and high-quality SNPs amplicon sequencing, our in-house approach enabled sensitive detection, quantification and classification of ON-target LOH as either deletion-based or copy-neutral. Indeed, in case of LOH, heterozygous SNPs are lost and are easily detected by sequence modification.

This robust strategy enables high-resolution, and accurate longitudinal monitoring of editing-associated chromosomal instability in clinically relevant cells, both *in vitro* and *in vivo*. It provides precise detection, description and mapping of ON-target genotoxicity at the single cell level. Our approach is more accurate than previous genotoxicity assays (CGH/SNP-arrays, optical genome mapping, FISH, PCR…) because (i) it relies on single cell analysis, more sensitive than bulk analyse, and because ii) it focuses on SNP in DNA, rather than scRNAseq^8, 12^, highlighting heterogeneity of outcomes (kilobase-and megabase-scale) and revealing frequent copy-neutral LOHs. Very recently, the same approach, scDNAseq has been also applied to study OFF-target genotoxicity and associated translocations^25^.

This approach was applied to human primary cells, fibroblasts and HSPCs, with editing performed at multiple loci on chromosomes 10 and 11. The theoretical sensitivity of detection of genotoxic events is 0.1%. To ensure high confidence and filter for technical artifacts, we defined a genotoxic event as a recurrent SNP profile observed in at least four individual cells. With this criteria, to reach 0.1% sensitivity, detecting a rare LOH event requires high-quality sequencing of 4,000 cells. In practice, by loading 75,000 to 100,000 cells in the Tapestri platform, we obtained around well-sequenced 3,000 fibroblasts and well-sequenced 1,000 HSPCs, leading to a sensitivity from 0.13 to 0.3%. This cell throughput is suitable for clinical applications.

scSNP-DNAseq generates a lot of data. We compared two bioinformatic pipelines to detect LOH: i) considering all the amplicons containing SNPs (global UMAP method Fig. 2), or ii) focusing on a set of five discriminant high-quality SNPs amplicons (SNP panel Fig. 3). We demonstrated that the second method, based on 5 discriminant SNPs is more performant than the global SNP approach.

Using this method, we identified heterogenous ON-target genotoxic outcomes reflecting the implication of several DSB repair pathways, and their dynamic time-course. Notably, genotoxicity in HSPCs was more severe when targeting *HBG1/2p* compared to *UROS*, showing higher frequency of both telomeric and interstitial LOHs, predominantly associated with deletions. This difference is likely due to DSBs occurring in two homologous sequences^24^. In a previous study, targeting another target in *HBG1/2p* in HSPCs^26^, we subcloned manually the edited cells and observed at day 30 large CN-LOHs after long-term amplification, associated with imprinting defects in *H19/IGF2* center^7^. Here, early analysis (day 4) revealed, for the first time in HSPCs, kilobase/megabase-scale deletions (5.2Mb encompassing 169 genes) in addition to CN-LOH in HSPCs. This contrast highlights the dynamic evolution of genotoxicity profile in HSPCs and underscores the need of longitudinal monitoring. In this context, scSNP-DNAseq emerges as a powerful and innovative tool for early genotoxicity detection and monitoring.

Understanding CRISPR-Cas9 ON-target genotoxicity is essential to anticipate and prevent potential risks. Spontaneous LOHs occur infrequently in human cells (<0.01%)^27^. The high-frequency of *in vitro* CN/CL-LOH starting from the cut site, interstitial or telomeric described in this work, is unlikely to be a random process. Diversity of genomic outcomes highlight the complexity of DSB repair. After DSB, NHEJ and homologous repair (HR) are the main mechanisms that resolve DSBs. In diploid cells, HR can result in interstitial copy-neutral LOH, either without or with cross-over. Alternatively, synthesis-dependent strand annealing SDSA (gene conversion) may create small interstitial copy-neutral LOHs. In case of chromosome truncations, the terminal chromosome-arm is lost by segregation errors during mitosis, inducing LOH associated with an extra-large deletion^28^. In primary cells lacking telomerase activity, the loss of telomeric regions is particularly detrimental. However, recently, *Jasin et al* reported that neo-telomere can be added^29^. If a neo-telemere cannot be added, we hypothesize that such events could be rescued by a secondary duplication of the remaining allele through break-induced replication (BIR), resulting in a CN-LOH.

When CRISPR is used for gene knockout, the error-prone NHEJ serves well. However, for precise genome editing, NHEJ can hinder the HDR, thereby reducing editing accuracy. To enhance HDR rate, DNA-repair modulators, commonly named HDR-boosters, have been proposed. We assessed the genotoxic risk associated with XL413^30^ and DNA-PKcs inhibitors (NU7441 and AZD7648), known to increase HDR^31–33^ or targeted integration^34, 35^. Our orthogonal approach revealed that AZD7648 poses a significant risk, as it induces megabase-scale ON-target genotoxicity *in vitro*, including a threefold increase in telomeric LOH and micronuclei increases in two primary human cells, targeting distinct genomic *loci*. Notably, genotoxicity associated with AZD7648 was not detectable at day 2 but became detectable later, highlighting the importance of delayed analysis. This observation aligns with a preprint from the Turchiano lab, which reported that NHEJ inhibition delays DSB repair^36^ and are in favor of chromosomal missegregation^37^. We hypothesized that, if Cas9-induced chromosomal DSBs are not repaired prior to mitosis, the resulting acentric telomeric fragments may missegregate, forming micronuclei that can drive chromosomal instability^9^. By blocking rapid NHEJ-mediated chromosome end-joining at the DSBs, DNA-PKcs inhibition likely increases the persistence of unrepaired DSBs, disrupting proper end pairing and elevating the risk of chromosomal truncations and long-term extra-large LOH. Our results are in accordance with recent papers: i) Corn *et al.*^18^ demonstrated AZD7648-induced genotoxicity (translocations and kilobase-scale deletions in HSPCs), ii) Jasin *et al.*^30^ observed extra-large LOHs (> 50Mb) following PolQ/DNA-PKcs inhibition in mouse embryonic stem cells and iii) Kosicki et al.^38^ reported kilobase-scale deletions in NHEJ-deficient mouse ES cells.

We now show that AZD7648 also increases micronuclei and megabase-scale CN-LOH events, which remain stable from day 4 to day 14 in HSPCs. These LOHs persisted for up to two weeks *in vitro* in HSPCs and for up to three weeks in fibroblasts, without apparent selective disadvantage. Notably, such AZD7648 genotoxicity can induce a bias in editing outcome analysis based on short-read amplicon sequencing^39^. This genotoxicity linked to DNA repair pathway modulation by drugs, could be expected regarding their use in oncology to sensitize cancer cells to chemo/radiotherapies/PARP inhibitors. These treatments are known to induce a catastrophic genomic instability ^16, 40–42^, demonstrating the critical role of NHEJ as a conservative mechanism that safeguards chromosomal integrity^43^.

Where millions of edited cells are transplanted, even rare genotoxic events become a major safety issue. While recent studies^8, 12, 19^ have shown that aneuploid CAR-T cells harboring chromosomal rearrangements are gradually lost during culture or post-graft in mice and patients, it remains unclear whether HSPCs behave similarly or persist *in vivo.* Importantly, in our study, after editing HSPCs at the *HBG1/2p* loci, we found no evidence of rearranged edited cells persisting *in vivo* after 3-months engraftment in NSG mice, even when editing was performed with genotoxic AZD7648. DNA-PKcs is known to also function beyond NHEJ, including ribosomal RNA processing and hematopoiesis. In mice, kinase-dead DNA-PKcs combined with *TP53* loss leads to hemopathies^44^. Here, transient DNA-PKcs inhibition prior transplantation did not impair HSPC engraftment or long-term hematopoietic differentiation. These new data suggest that AZD7648-induced genotoxicity in chr11p can be lost *in vivo* but has to be carefully monitored. Very little is known about the fate of rearranged cells post-grafting. It is well-known that missegregated chromosomes often suffer during cytokinesis, triggering a DNA DSB response in daughter cells, involving ATM, Chk2, and p53, potentially leading to cell death^45^. Further studies are needed to decipher the precise time-course of disappearance of these unwanted genomic by-products. It will be essential to determine if their *in vivo* loss results from i) impaired graft capability, ii) *in vivo* selective disadvantage of edited cells with genomic rearrangement, iii) intrinsic protection of quiescent HSPCs ON-target genotoxicity, in accordance with G0/G1 protection. Whatever the underlying mechanisms, these *in vivo* results provide encouraging evidence supporting the safety of CRISPR-Cas9 nuclease protocols at least for the conditions tested.

Although we observed the disappearance of genotoxic events at long-term following *HBG1/2p* targeting in Chr11p, we cannot exclude that it is *locus* specific and that other chromosomal rearrangement could persist and potentially confer a selective advantage *in vivo*. Prevention of the initial occurrence of ON-target chromosomal genotoxicity is the best approach to propose safe CRISPR gene therapy. The best way to avoid genotoxicity is not to hope for its disappearance over time but rather to prevent its initial appearance. In this study, we also demonstrated the accuracy of scSNP-DNAseq to evaluate and refine editing protocols, to mitigate ON-target genotoxicity. For the first time, we show that palbociclib, a CDK4/6 inhibitor, drastically lowers CRISPR-induced ON-target genotoxicity in HSPCs. We hypothesized that editing non-dividing HSPCs could bypass unintended genotoxicities associated with DSBs in dividing cells. Our data demonstrate that palbociclib results in less MN and in a 7-fold reduction in telomeric megabase-scale LOH, without impairing neither gene knockout efficiency by NHEJ nor cell graft capability. This preventing effect is in accordance with previous findings indicating that i) micronuclei and long deletions predominantly occur in dividing cells^46–48^, that ii) gene editing of quiescents HSPCs (without *ex vivo* culture) avoids genotoxicity^20^, and iii) gene editing of non-proliferating CAR-T editing is safer compared to proliferating T-cell^12^.

We proposed that G0/G1 arrest during editing could impair genotoxicity by promoting end-joining after DSB. When palbociclib was combined with DNA-PKcs inhibition, to block NHEJ, genotoxicity mitigation was impaired, despite G0/G1 arrest, supporting the hypothesis that palbociclib may reinforce NHEJ repair pathway to preserve genomic stability. Altogether, these data highlight the central role of cell cycle in the regulation of ON-target genotoxicity, and underscore the utility of scDNA-seq for screening and optimizing safer genome editing protocols. In a future clinical projection, we evaluated palbociclib impact on long-term HSPCs function. Unexpectedly, palbociclib enhances HSPCs stemness phenotype and graft capability. Taken together, these results demonstrate the potential of palbociclib as a valuable strategy to mitigate CRISPR-Cas9 nuclease ON-target genotoxic risk for gene therapy.

Alternatively, new DSB-less CRISPR tools (e.g. base, prime and click editors^49–53^) have been proposed to minimize genotoxicity. However, recent studies reported unintended ON-target genotoxicity following base/prime editing in HSPCs^54, 55^. This suggests that scSNP-DNAseq, could also become an essential platform to assess the safety of the newer editing approaches.

## Limitations of the study

Our custom panel enables sensitive detection and mapping of ON-target genotoxicity, particularly kilobase/megabase-scale rearrangements. This panel was designed to study two chromosomes (10 and 11), with approximately 400 amplicons containing SNPs analyzed at single cell level. However, it does not provide a whole genome picture. Another team proposes a genome-wide approach, using a custom panel spanning all 24 chromosomes with one amplicon each 10Mb. They detect whole chromosome aneuploidies by CNV analysis (preprint ^56^). However, this method lacks the resolution detection of aneuploidies in shorter regions, and copy-neutral genomic events. Nowadays, it is required to make a trade-off between sensitivity and genomic breadth analysis. Further optimizations of such panels combining local and whole genome coverage will allow the precise global analysis of genotoxicity. In this work, for bioinformatic analysis, we demonstrated that a focus approach using a small number of high quality SNPs (five discriminant SNPs) outperforms global SNP analysis (UMAP) to detect megabasic et kilobasic LOH. Areas for future panel optimization include: i) SNP pre-identification to choose only amplicons with SNPs truly present in the sample, ii) improved amplicon pre-selection based on high PCR amplification efficiency (high total read counts) and iii) expanding the panel size (current Tapestri technology allows up to 800 amplicons per cell, enabling broader coverage for both ON- and OFF-target genotoxicity detection, as proposed by Kogenaru et al.^23^).

Another limitation is cost. The microfluidics, library prep, and sequencing components of this technology are currently expensive, making it less accessible for routine use in academic laboratories. However, costs could be reduced by sample multiplexing, and this approach could be implemented in clinical gene therapy workflows for pre- and post-graft quality control. In this context, the cost of safety monitoring must be weighed against the high expense of cell and gene therapies themselves.

In conclusion, our study demonstrates that multiplex single-cell approaches, and in particular custom SNP-based scDNA-seq, is a robust and sensitive tool that outperforms current methods, including scRNA-seq, for detecting at the single-cell level in primary cells *in vitro* and *in vivo* copy-loss and copy-neutral events. We identified frequent ON-target genotoxicity and chromosomal rearrangements post-editing, especially with AZD7648 and demonstrated that they can spontaneously disappear *in vivo* (targeting *HBG1/2p)* or be mitigated by palbociclib. We propose micronuclei (automatic quantification) and scSNP-DNAseq integration as a quality control step in (pre)clinical genome editing workflows. It could be implemented both before transplantation of CRISPR-Cas9-edited cells and post-graft, to monitor the long-term behavior of rearranged cells.

## Acknowledgements

Tapestri equipments from Mission Bio were acquired at Gustave Roussy Institute thanks to MyProbe ANR-17-RHUS-0008. This work was supported by the French national research agency (ANR) (grants ANR-21-CE18-0002 and ANR-PRME-23-CE52 and by la Fondation pour la Recherche Médicale (FRM, grant EQU202403018062). C.B. was supported by funding from INSERM (Poste accueil INSERM). C.T. is supported by AFM telethon (#24581). We thank Ivan Lukic and Stephane Collinet from Mission Bio for their advice and technical support for Mission Bio software. We thank Zoran Ivanovic, research director at Etablissement Francais du Sang (Aquitaine, France), for his valuable advice concerning hematopoietic stem cell *in vivo* assays. We thank the FACSility cytometry platform at Bordeaux University (TBMCore, Bordeaux, France). We thank INSERM transfert for valorization assistance. We thank Sandrine Hamon, Patrice Dutrinus and Stephanie Lannelongue for administrative assistance and financial management.

## Author contribution

**V.M, S.F., C.B., J.B., A.B., F.M.G**. design, analysis of data and drafting of the manuscript. **I.L-G, S.F., M.R**. **C.B.**: cell culture, CRISPR editing, FACS experiments. **C.B.**, **C.T., V.M.**: nanopore sequencing, RTqPCR and bioinformatic analysis; **N.D**., **M.F.**: scDNA-seq experiments. **A.P**.: cell cycle protocol development. **V.M., J.B.**: scDNA seq bioinformatics analysis; **J.T**.**, J.B.**: FISH experiments. **P.B.G., C.D., J.B.** : animal study design and experiments. **S.D**., **J.B.**, **D.C**.: helpful discussion and revision of the manuscript. **A.B., F.M.G**.: supervision, funding acquisition, analyzing data, writing the manuscript, final approval of the manuscript. All authors edited and approved the final manuscript.

## Disclosures

S.F., A.B., and F.M.G declared a patent application EP23305760 filed on May 13th 2023 and claims the use of palbociclib to prevent genotoxicity induced by nuclease. The remaining authors declare no competing interests.

## Methods

### Ethics statement

Our research complies with all relevant ethical regulations. Cord blood Human CD34^+^ HSPCs were obtained in collaboration with EFS Nouvelle Aquitaine-Limousin (24SPL 02 agreement for the cessation of between EFS and Bordeaux University), according to the hospital’s ethical institutional review board and with the mother’s informed consent. Prior to donating peripheral blood, each donor read an informative document and completed a questionnaire allowing the possibility to be opposed to the use of discarded cells for research.

### Cell culture

Human foreskin fibroblasts immortalized with hTERT (hFFs) were from ATCC® (CRL 4001, BJ-5ta). They were partially inactivated for *UROS* using a ribonucleoprotein (RNP) made of Cas9 protein complexed with a gRNA targeting *UROS* exon 4^17^. hFFs were maintained in DMEM, high-glucose (4.5 g.L-1), L-Glutamine (1 g.L-1) and pyruvate (Gibco® by Life-technologies^TM^, Carlsabad, CA, USA, catalog n°31966047) supplemented with 10% fetal bovine serum FBS (Eurobio, Les Ulis, France, Catalog #CVFSVF00-01), 100 U/mL penicillin and 100μg/mL streptomycin (Gibco® by Life-technologies^TM^, Carlsabad, CA, USA; catalog #15070063), 10 µg/mL ciprofloxacin (Biogaran™, Colombes, France) and 0.5 µg/mL amphotericin B (Sigma-Aldrich®, Saint Louis, MO, USA, catalog #A2942).

Human CD34^+^ HSPCs were isolated from the cord blood. Briefly, mononuclear cells were isolated by Ficoll gradients. HSPCs were purified according to the manufacturer’s instructions (hCD34-Positive Selection kit II, from Stem Cell Technologies, Vancouver, BC, Canada, catalog #17865) and purity was analyzed by flow cytometry using phycoerythrin (PE)-conjugated anti-CD34 antibody (clone 561, Biolegend,San Diego, CA, USA). The CD34^+^ cell purity was> 90%.Cryopreserved HSPCs were thawed and cultured in expansion medium consisting in StemSpan SFEM (Stem Cell Technologies, catalog #09600) supplemented with Flt3-L (100 ng/mL, catalog #GMP300-19-50UG), SCF (100 ng/mL, catalog #GMP300-07-50UG), hTPO (100 ng/mL, catalog #GMP300-18-50UG), (all from Peprotech, Cranbury, NJ, USA), 100 U/mL penicillin, and 100 μg/mL streptomycin (Gibco® by Life-technologies^TM^, Carlsabad, CA, USA, catalog n°15070063), 10 µg/mL ciprofloxacin (Biogaran™, Colombes, France) and 0.5 µg/mL amphotericin B (Sigma-Aldrich®, Saint Louis, MO, USA, catalog n°A2942).

All cells were grown at 37°C with 5% CO2 levels. All cell lines were tested for mycoplasma.

To increase HDR, we added 3 “HDR boosters” in the medium for editing : NU7441 (2.5 µM, 18 h before transfection and for 2 days after transfection, from Selleckchem, Houston, US, catalog #S2638), AZD7648 (1 µM, 18 h before transfection and for 2 days after transfection, from MedChemExpress, Monmouth Junction, New Jersey, US, catalog #HY-111783) or XL413 (20 µM, for 2 days after transfection, no cell exposure performed before transfection, from Selleckchem, Houston, US, catalog #S7547). Dimethyl sulfoxide (DMSO, from Sigma-Aldrich®, Saint Louis, MO, USA, catalog #D2438), used to dissolve these compounds was used as a control. To prevent genotoxicity, cells were synchronized in G0/G1 phase by incubation with palbociclib (1 µM, Sigma Aldrich, Saint Louis, MO, USA, catalog #PZ0383) 18 h before and 24h or 48h (for scDNA-seq) after RNP transfection.

### Editing tools

The crRNA for the *ABRAXAS2* and *UROS* targets (Ch10q) was designed using CHOP-CHOP software (chopchop.cbu.uib.no) (*ABRAXAS2* 5’-GAGATCCTCCCACTCGATGG-3’ and *UROS* exon 4: 5’- GGAAGCAGCAGAGTTATGTT-3’). The crRNA chosen for editing of the *HBG1* and *HBG2* promoters (Chr11p) is an already published gRNA (gRNA-68)^24^. The different components of the CRISPR-Cas9 system were combined to form a RNP (all ordered at IDT, Integrated DNA Technologies, Coralville, IA, USA). For this, 17 µg of Alt-R® S.p. HiFi Cas9 Nuclease V3 (abbreviated hereafter as Cas9, Integrated DNA Technologies, Coralville, IA, USA, catalog #1081059) and 6.5 µM of sgRNA, were mixed and then incubated for 10-20 minutes at room temperature before electroporation. For HDR booster experiments, 6.5 µM of ssODN template was added (*ABRAXAS2* 5’- TCTGCTTCTTCATCCTAGTTCACTCACTTTGCCCCCAGCTGCCT**CCATCGAGTGGGAGG*AA*GAGCTCGA**CAGAG CCTGTTCACTCCTAGtcatttatgcaacagacatcaatggaaacat-3’). Finally, 3.9 μM of Alt-R Cas9 Electroporation Enhancer solution (Integrated DNA Technologies, Coralville, IA, USA, catalog #1075916) was added to the RNP complex to improve electroporation efficiency.

Cells were transfected by electroporation using the Nucleofector 4D AMAXA electroporation system (Lonza®, Bale, Switzerland). In brief, 2.10^5^ to 3.10^5^ depending on the cell type (HSPCs and hFFs) were resuspended in P3 Primary Cell Line 4D-Nucleofector® (Lonza, Bale, Switzerland, catalog #V4XP-3024) and added to the RNP complex. Then cells were nucleofected using DO-100 and CZ-167 programs respectively.

### Editing efficiency

Genomic DNA of edited cells, and their associated controls, was extracted using Nucleospin® Tissue (Macherey-Nagel®, Düren, Germany, catalog #740952.250) according to the manufacturer’s protocol. The genomic region flanking the expected cut-site was amplified by PCR (HotStarTaq Plus DNA polymerase, Qiagen®, Venlo, Netherlands, catalog #203605) with adequate primers (*ABRAXAS2* targeting F: 5’-CCTGGCTCACTCACTCTGTG-3’, R: 5’-AGGCCAGTTCAAACGCTCTT-3’; *UROS* targeting F: 5’- TAGTTCCAGGCACATAGTAAGCAC-3’, R: 5’-TCCCAAGGCAGAGTCTGTGA-3’; *HBG1/2* promoters targeting F: 5’- ACGGCTGACAAAAGAAGTCCT -3’, R: 5’-AGCCTTGTCCTCCTCTGTGA-3’). PCR products were purified with Nucleospin® Gel and PCR Clean-up (Macherey-Nagel®, Düren, Germany, catalog #740609.50S). Sanger sequencing was done on purified PCR products and sequenced by LIGHTRUN (GATC Biotech, Konstanz, Germany). Sanger sequencing data were analyzed using ICE v2 CRISPR Analysis tool (ICE) software (Synthego, Redwood City, CA, USA) and DecodR v3^57^. Purified PCR products from unedited cells were used as control chromatograms.

To validate CRISPR-Cas9 efficiency at HBG1/2 locus in HSPCs, we used Nanopore NGS. Briefly, 7 days after editing, genomic DNA was extracted from cell pellets using Nucleospin® Tissue (Macherey-Nagel®, Düren, Germany, catalog #740952.250) according to the manufacturer’s protocol. The genomic region flanking the edited site was amplified by Long Range PCR PCR (LA Taq, Takara Bio, Kusatsu, Japan), with adequate primers. Sequencing library was prepared using Native Barcoding Kit 96 V14 (Oxford Nanopore Technologies, Oxford, United-Kingdom, catalog #SQK-NBD114.96) according to the manufacturer’s protocol. Sequencing was then performed using a MinION Mk1B device and a Flow Cell R10.4.1 (Oxford Nanopore Technologies, Oxford, United-Kingdom, catalog #FLO-MIN114) to reach at least 3000 qualify-reads per sample (Q score > 9). After sequencing, super-accurate basecalling was performed using MinKNOW (v24.02.6) and fastq were aligned on human reference genome GrCH38 (GCA_0000001405.15) using a home made Bash (v5.1.16) script based on Epi2Me (v5.1.9) wf-alignement pipeline (v1.1.2). Then, BAM were analyzed using a homemade bioinformatic pipelines in Python (v3.13.2) using pysam (0.22.0). Briefly, it classifies reads into categories (WT, small inDels, 5 kb deletions, truncated etc…) based on CIGAR string interpretation. Results are summarized in .csv tables using pandas (v2.3.1). All scripts and documentation are available at: https://github.com/BioGO-BRIC/HBG1-2---ONT-analysis

### Micronuclei analysis by cytometry

The MicroFlow In Vitro-250/50 Kit (Litron Laboratories, Rochester, NY, USA, catalog #INVITRO250/50) was used for all MN assays, following the manufacturer’s instructions. Briefly, cells were stained and lysed following the protocol described in the kit. Following harvest, a photo-activated dye Dye 1, Ethidium monoazide (EMA) was used to stain apoptotic and necrotic cells then the cells were washed. Subsequently, healthy chromatin was stained with the nucleic acid Dye 2, SYTOX Green. Cells could be stored up to 24 hours at ambient temperature, protected from light, or refrigerated for up to 72 hours, prior to analysis. Using a BD Accuri C6 Plus (BD Biosciences, Franklin Lakes, NJ, USA), flow cytometric analysis distinguished 10.000 to 20.000 real MN (SYTOX Green only) from nuclei, based on their weaker DNA-associated fluorescence signal.

### MN analysis by nuclear fluorescent imaging

Cytogenetic examination was carried out on interphase nuclei on HSPCs after fixation by a solution of 3:1 methanol/glacial acetic acid fixative. DAPI (Sigma Aldrich, Saint Louis, MO, USA, catalog #MBD0015) staining was performed onto a microscopic slide and MN examination was evaluated under a fluorescent microscope Leica DM5500 B (Leica microsystem, Wetzlar, Germany) with the 63x objective lens (i.e. 630x magnification). The number of cells visualized for each condition is indicated in the legend of each figure. Blind count of micronuclei with a medical cytogeneticist (J.T. in author list).

### LOH analysis by FAMReD system (cytometry)

FAMReD (Fluorescence-Assisted Megabase-scale Rearrangements Detection) is a cytometry-based system to detect induced LOHs in Chr10q with or without *TP53* inactivation^17^. On day 15 post-editing of hFFs *UROS*^+/-^, 0.3 mM of 5-ALA (Sigma Aldrich, Saint Louis, MO, USA, catalog #A7793) were added to culture media. After 16 h of exposure (overnight), cells were washed twice with DPBS 1X (Gibco, by Life technologies, Carlsabad, CA, USA, catalog #14190144) and put back in fresh media. Upon LOH, loss of *UROS* can be detected by the appearance of fluorescence due to porphyrin accumulation. Following 8 hours of clearance, fluorescent cells were quantified by flow cytometry using a BD Accuri C6 Plus (BD Biosciences, Franklin Lakes, NJ, USA). UV-sensitive porphyrins were excited at 488 nm and the emitted wavelength was approximately 667 nm, detected by the PE-Cy5A PMT channel.FL-1 is a control green-fluorescent channel used to exclude auto-fluorescent cells. The system only detects 50% of the events, when the cells become UROS ^-/-^. The true LOH rate is double.

### Cell cycle

To evaluate impact of palbociclib and HDR boosters on hFFs and HSPCs cell cycle, cells were harvested and washed with DPBS (Gibco® by Life-technologiesTM, Carlsabad, CA, USA, catalog #14190144) with 2% FBS (Eurobio, Les Ulis, France, catalog #CVFSVF00-01) and EDTA 2mM (Éthylènediamine-tétraacétate disodium dihydrate, Euromedex, Souffelweyersheim, France, catalog #EU0007). Cells were fixed and permeabilized using the True-Nuclear Transcription Factor Buffer Set following the manufacturer’s instructions. True-nuclear Fixation buffer (1X, from True-Nuclear Transcription Factor Buffer Set, 424401, BioLegend, San Diego, California, USA, catalog #424401) was added and incubated for 45 min in the dark at ambient temperature. Subsequently, a true-nuclear 1X Perm Buffer (from the same set, BioLegend, San Diego, California, USA, catalog #424401) was used twice to wash. Cells were then stained with a PE mouse anti-Ki67 antibody (BD Biosciences, Franklin Lakes, NJ, USA, catalog #556027) for 30 min in the dark at ambient temperature and washed with true-nuclear 1X Perm Buffer (from the same set, BioLegend, San Diego, California, USA, catalog n°424401). Pellet was harvested in DPBS with FBS 2% and EDTA 2mM and resuspended in a 2µg/mL Hoechst solution (bisBenzimide H 33342 trihydrochloride, Sigma-Aldrich®, Saint Louis, MO, USA, catalog B2261). Samples were stored at +4°C overnight before analysis performed on a BD LSRFortessa™ Cell Analyzer (BD Biosciences, Franklin Lakes, NJ, USA). Data were analyzed with BD FACSDiva™ Software (BD Biosciences, Franklin Lakes, NJ, USA).

### Design of custom Tapestri panel for sequencing

Panel comprises 403 probes mainly across the 10 & 11 human autosomes (Extended Data Table 1). To identify candidate target regions for the panel, we used the dbSNP (NCBI dbSNP Build 155, hg19) and considered only variant SNPs with a major allele frequency between >0.5 and <0.7 without discrepancy on certain ancestry group (eg global MAF = 0,5 and <0,5 in African and Caucasian ancestry in GnomAd v4). *In Silico* validation of the SNP design was assessed by testing it on the CORIELL (GIAB consortium, sample NA12878, HG001) using bcftools (v1.18). The objective of >30% informative SNPs was largely achieved with 63%. All candidate SNPs were submitted to the Tapestri Panel Designer to generate a panel design and ensured that the designed probes targeted the candidate SNPs and had similar GC contents. Support for the custom panel design and synthesis of the panel was provided by Mission Bio using v3 Tapestri Chemistry (Mission Bio, South San Francisco, CA, USA).

### Single-Cell DNA-seq

Single-cell DNA-seq was performed using the Tapestri platform (Mission Bio, South San Francisco, CA, USA) according to the manufacturer’s specifications with kit v3. Briefly, cryopreserved human cells were gently thawed, washed, and quantified using a Countess II cell counter (ThermoFisher Scientific, Waltham, MA, USA). The cells were then diluted to a concentration of 2,000 to 3,000 cells per μL in the cell buffer. Next, 35 μL of cell suspension was loaded onto a microfluidics cartridge and cells were encapsulated on the Tapestri instrument followed by cell lysis with protease digestion followed by heat inactivation using a thermal cycler. The cell lysate was reintroduced into the cartridge and processed such that each cell possessed a unique molecular barcode. Amplification of the custom targeted DNA regions was carried out by incubating the barcoded DNA emulsions in a thermocycler following Mission Bio’s specifications. Emulsions were broken, DNA digested and purified with AMPure XP beads (Beckman Coulter, Brea, CA, USA, catalog #A63881). The beads were pelleted and washed and DNA libraries were generated. Final libraries were purified with AMPure XP beads. All libraries were sized and quantified using an Agilent Fragment Analyzer and pooled for sequencing on an Illumina NovaSeq6000 S1 flowcell with 2 x 150bp at Gustave Roussy. A summary of metrics (mean reads/cell/amplicon, reads mapped to target, dropout rate and amplicon > 10X) are provided in extended table 4. Data for each run and sample are in raw data (Zenodo).

### Tapestri Single Cell DNA analysis

Single-cell DNA sequencing data were obtained from primary cells and analyzed using the Tapestri Mosaic pipeline and customized versions of Mission Bio’s available jupyter workflows (Mission Bio, San Francisco, CA, USA). Quality control steps included the removal of low-quality or missing genotype cells as per the pipeline’s automated filters. Variants were annotated using the default settings provided by the manufacturer in the Tapestri Mosaic suit.

For HSPCs edited in *HBG1/2* promoters, clustering method using principal component analysis (PCA) and UMAP, based on 88 heterozygous SNPs on chromosome 11 (Chr11) was performed. The optimal number of principal components (PCs) was determined through an elbow plot analysis, identifying the inflection point where the explained variance ratio begins to plateau. This transition, occurring around 8 PCs (determined via visual inspection), represents the balance between maximizing data variance retention and minimizing noise contribution. UMAP were then generated using the selected PCs as input, employing the Euclidean distance metric with default Tapestri Mosaic parameters. Clustering was carried out in an unsupervised manner using the *hdbscan* algorithm.

In case of selection of 5 high-quality SNPs, variant annotation manufacturer settings were not modified. LOHs analysis was performed by adjusting the max ADO score at 0.2 and the Minimum clone size at 0.01. Only heterozygous SNPs (0.40<AF<0.60) present in at least 60% of the cells were retained.

SNP profiles to cluster cells were retained if they contained at least 6 cells. The CNVs analysis was only performed in the LOH subtypes with at least 6 cells. The filters were set up to 50 for amplicon completeness, 10 for amplicon read depth and 10 for mean cell read depth. Copy number variations were inferred from single-cell amplicon read-depth data using the Mission Bio Mosaic pipeline. Ploidy was calculated for each cell and each amplicon by normalizing read counts against the mean of the “normal” cell population, which was designated as the diploid reference (considering ploidy 2 for normal cells, without LOH). LOH ploidy was calculated by averaging the ploidies obtained for amplicons encompassing the LOH.

### Proliferation, cell death, *in vitro* phenotype and stemness expression in HSPCs

Proliferation was assessed with sequential cell numeration following or not 36-hour exposure to Palbociclib. Cell numerations were performed using KOVA slides 10 counting chambers (Alltrista Plastics Ltd., USA; distributed by Labellians, France, Ref. 87144F) according to the manufacturer’s instructions.

Apoptosis is detected by staining the cells with anti-human APC Annexin V (#640920 Biolegend, San Diego, CA, USA) and propidium Iodide (#596552, Dutscher) solution followed by flow cytometry analysis. Cells that were propidium iodide (PI) negative and Annexin V negative are considered healthy, cells, PI negative and Annexin V positive cells are considered apoptotic, and cells that are positive to both PI and Annexin V considered necrotic.

To evaluate phenotype (stemness) of HSCPs, cells were stained with human FITC anti-CD38, and human PeCy7C anti-CD90 antibodies, from BD Biosciences (respectively #567148 and #561558, BD Biosciences,Franklin Lakes, NJ, USA) human PE anti-CD34 and human APC anti-CD133 from Biolegend (respectively #343506 and #372806, Biolegend, San Diego, CA, USA), and analyzed by cytometry using FACS Canto II from BD.

Stemness Hematopoietic gene expression is evaluated by RTqPCR. Total RNAs were extracted using Direct-zolTM RNA MiniPrep (Zymo Research, Irvine, California, USA) or the RNeasy Micro kit (Qiagen). mRNAs were reverse transcribed using the High-Capacity cDNA Reverse Transcription Kit (Applied Biosystems, Life Technologies Corporation, Carlsbad, California, USA). In all cases, qPCR was performed using Go Taq® qPCR Master Mix (Promega, Fitchburg, Wisconsin, USA) in a CFX96TM Real-Time System (Bio-Rad Laboratories, Hercules, California, USA). Primers were synthesized by Eurogentec (Liège, Belgium). Primers previously published^58^ were used.

### Hematopoietic engraftment in immunodeficient mice

The animal experiments were authorized by the Ethical Committee of Gironde (CEEA50, 2017/03/30), France, and Ministère de l’Enseignement Supérieur et de la Recherche (ref. APAFIS #35068-2022013110253304 v8) and are in agreement with French regulation—licenses granted to C.D. and P.B.G. through authorization n°33 07 10 from Préfecture de la Gironde and A33 12 054 from Ministère Français de l’Enseignement Supérieur et de la Recherche, respectively. NSG mice were obtained from the Jackson laboratory and bred in-house (animal facilities, University of Bordeaux). Female mice between 8-10 weeks old were used for all experiments. Prior to cell injection, mice were conditioned by two intra-peritoneal injections of 20 mg/kg busulfan (Busulfan; Fresenius Kabi), at 24h intervals. Two days after the second injection of busulfan, unmanipulated and treated hCD34^+^ cells were IV-injected into recipient mice under anesthesia (Isoflurane, 5% at initiation then 2.5%). Depending on our objectives, mice were sacrificed at different time points (weeks) after transplantation, and bone marrow (BM) was harvested from femurs. Total BM cells were labelled with BV421 anti-murine TER119 (BD, 740686), BV421 anti-murine CD71 (BD, 740667), BV421 anti-murine CD45 (BD, 563890) to exclude murine cells from the analysis. FITC anti-human CD45 (Beckman Coulter, A07782), PE anti-human CD19 (Beckman Coulter, A07782), and APC anti-human CD33 (Beckman Coulter, A07782) antibodies. Human vs. mouse chimerism was analyzed by flow cytometry with FACS Canto II.

#### Palbociclib effect

For Limiting dilution assay (LDA) with or without palbociclib, ten mice per group (cell dose) received 25, 50, 100, 500 or 3000 CD34^+^ cells non-treated or treated with palbociclib. Mice were sacrificed 17 weeks after transplantation and their bone marrow analyzed by flow cytometry as mentioned above. SRC frequency with 95% IC were determined using the extreme limiting dilution analysis (ELDA) webtool accessible at http://bioinf.wehi.edu.au/software/elda/. For secondary transplantation, to evaluate a potential effect of palbociclib on long-term SRC, 3000 non-treated or treated CD34^+^ cells were injected in primary mice (10 mice per group). 8 weeks after transplantation, mice were sacrificed and bone marrow (BM) was harvested from femurs. The total cells from one femur of each primary mouse was suspended in 20µL of StemSpan SFEM II medium (StemCell Technologies) and then injected directly in the bone marrow of one respective secondary recipient by intra-bone injection. Mice were sacrificed 17 weeks after transplantation and their bone marrow analyzed by flow cytometry as mentioned above.

For Fig. 4g, in order to test palbociclib function on engraftment, 5 mice per group received 10 000 CD34^+^ cells treated or not with palbociclib (n=2 independent experiments). For supplementary Fig. 5, 5 mice per group received 50000 cells, edited or not in *ABRAXAS2 by CRISPR-Cas9,* with or without palbociclib (n=2 independent experiments). Mice were sacrificed 17 weeks after transplantation and their bone marrow analyzed by flow cytometry as mentioned above.

#### long-term *in vivo* genotoxicity monitoring (with or without AZD7648)

To assess the potential genotoxicity of HBG1/2p-CRISPR editing on CD34^+^ cells, 400 000 CD34^+^ edited without or with AZD7648 were injected intravenously into recipient mice (2 mice per group). 90 days later, the mice were sacrificed and the whole bone marrow cells from femurs and tibias were harvested prior to human CD45^+^ cell selection by magnetic beads (Millteny biotech, hCD45 microbeads) and scSNP-DNAseq experiment.

### Statistical analysis and reproducibility

When possible, experiments were conducted at least 3 times independently. No statistical method was used to predetermine sample size. No data were excluded from the analyses. The experiments were not randomized. DAPI analysis was performed blindly. Statistical significance was inferred when necessary. Exact distinct and independent experiments size is indicated in each legend (*n*). Graph Pad Prism 10 software (GraphPad, Boston, MA, USA) was used for statistical analysis. Results are presented as mean ± SD, frequencies or mean ± SEM (indicated in each figure legend). The parametric *T* test (two-sided) was used when distribution was Gaussian/normal (Shapiro–Wilk test). The non-parametric Mann–Whitney test (one-sided) was used to compare two groups when distribution was not normal. One-way ANOVA (two-sided), complemented with the unprotected Fisher’s Least Significant Difference test, was used to compare more than two groups. Percentages of LOHs in HSPCs were compared by the Chi-square test.

## Data availability

Source data are provided with this paper. The source data generated in this study have been deposited in the ZENODO database https://doi.org/10.5281/zenodo.16638384. FASTQ files of Single cell DNA sequencing and Long read nanopore Minion can be provided on demand.

---

**Supplementary Fig. 1:**
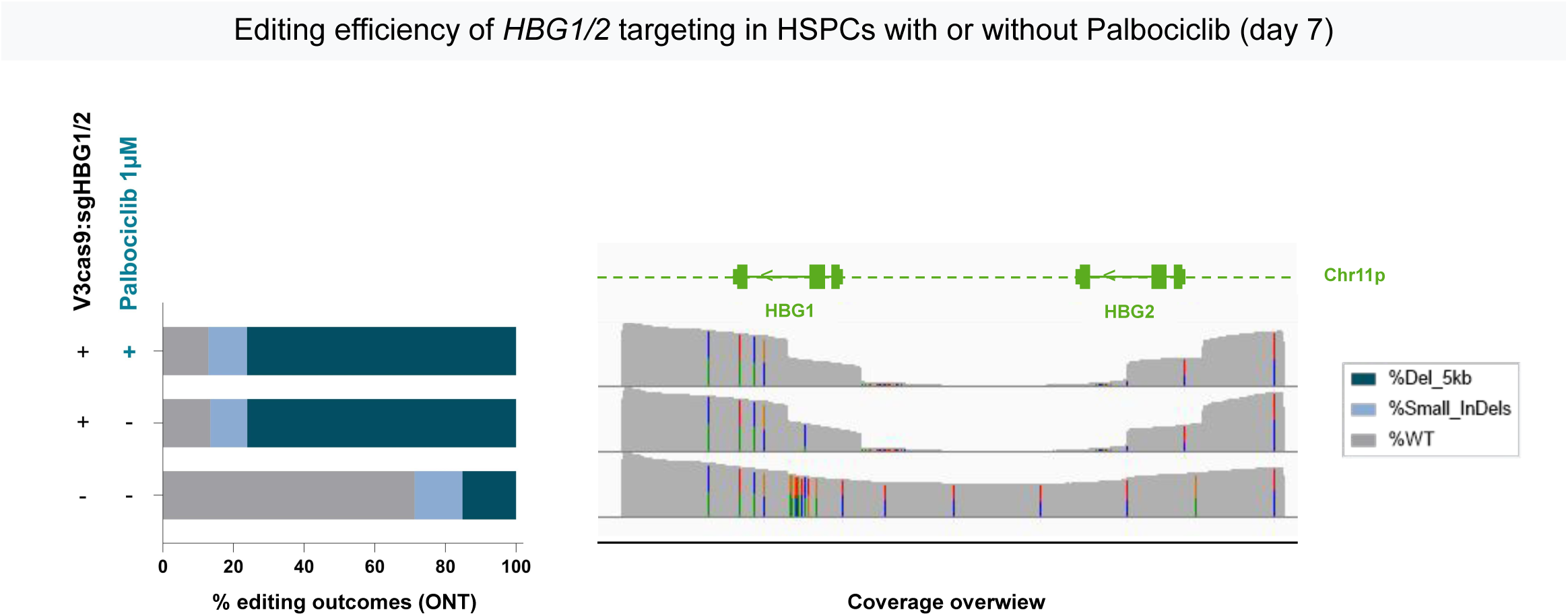
Editing efficiency measured by long-range PCR (10 kb) and Nanopore sequencing for *HBG1/2* editing. **Left panel**, histogram of genomic events after *HBG1/2* promoter targeting by CRISPR-Cas9 nuclease in HSPCs. Grey = WT sequence; light blue = indels, mutation or sequencing error; dark blue = 5Kb-deletion in HSPCs non edited, edited, and edited with palbociclib. **Right panel,** coverage overview of NGS by Nanopore visualized with IGV.

**Supplementary Fig 2:**
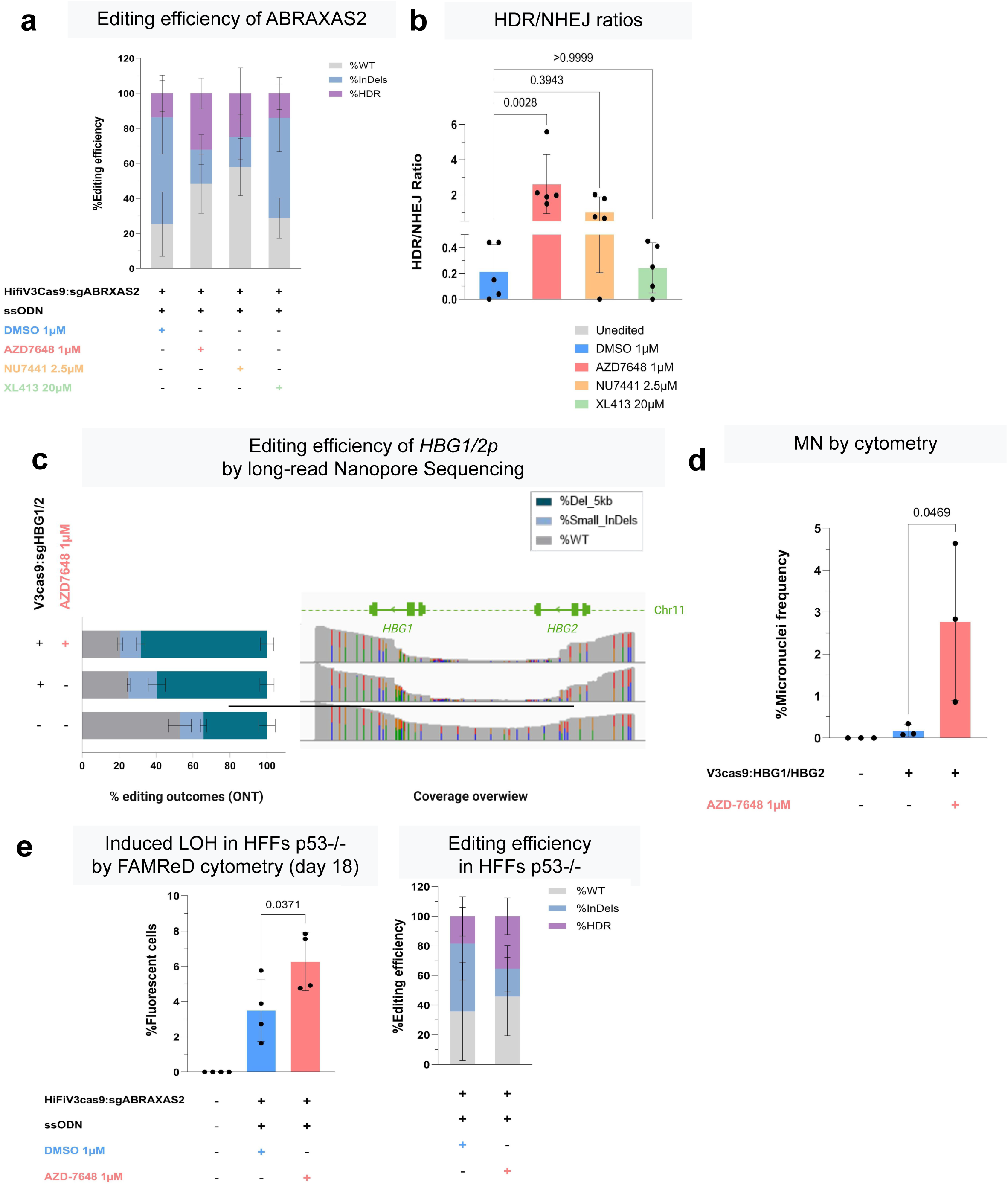
Genotoxic effect of AZ7648 in hFFs. **a,** Editing efficiency of *ABRAXAS2* editing in presence of HDR boosters, measured by PCR and DECODR 7 days after editing (Mean ± SD). **b**, HDR/NHEJ ratio calculation by DECODR software (Mean ± SD, n=5 independent experiments, One-way ANOVA test). **c**, Editing efficiency of *HBG1/2p* editing 7 days after transfection, by long-range PCR and Nanopore sequencing. **d**, Micronuclei quantification by cytometry (day 2 after *HBG1/2p* editing) with or without AZD764 (Mean ± SD, n=3 independent experiments, One-way ANOVA). e, **left panel**: megabase-scale LOH (fluorescent cells) quantification by FAMReD after *ABRAXAS2* targeting in p53-deficient hFFs (Mean ± SD, n=4 independent experiments, One-way ANOVA). **right panel**: Editing efficiency of *ABRAXAS2* editing in p53-deficient hFFs, measured by PCR,

**Supplementary Fig 3:**
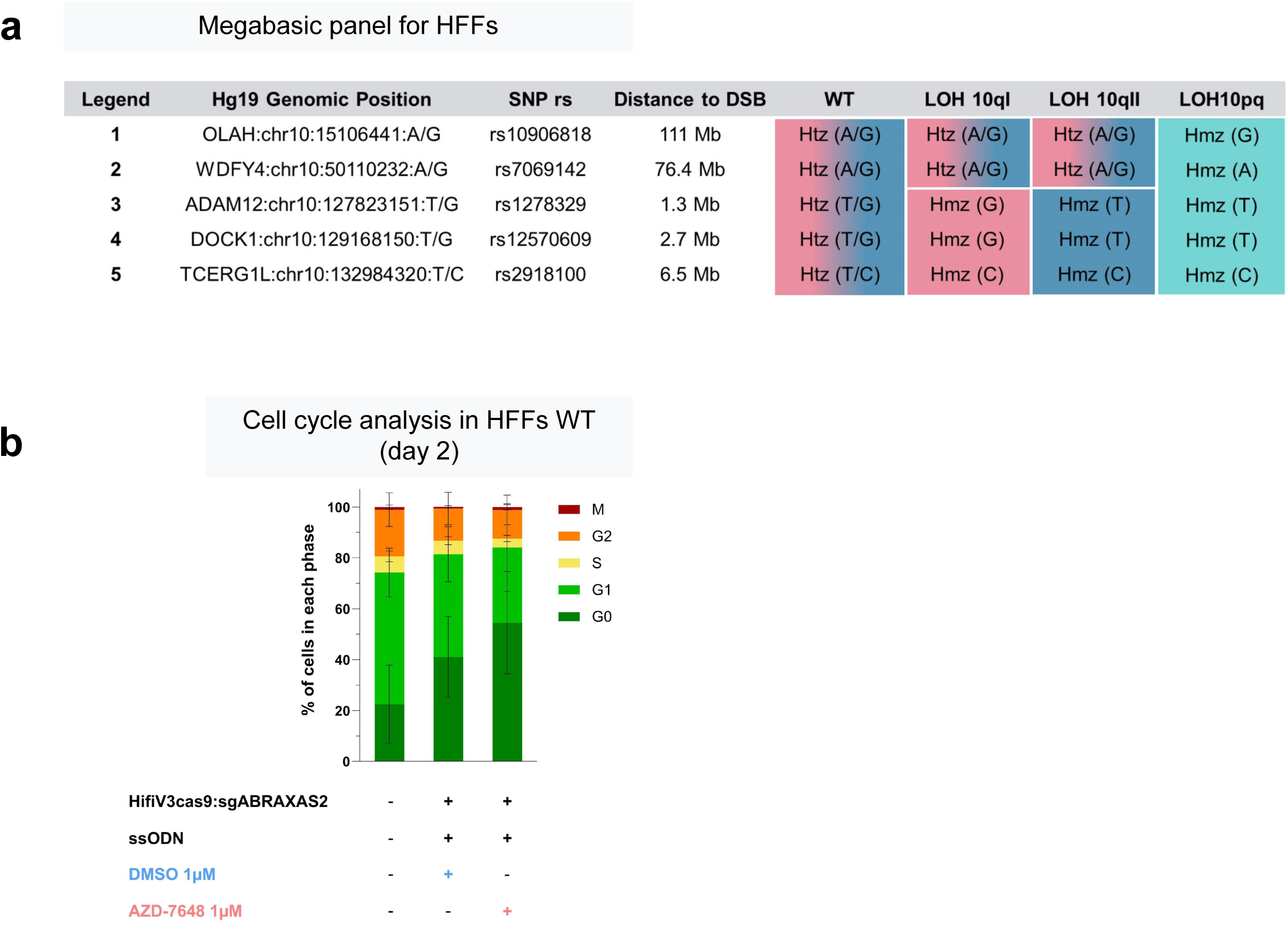
AZD7648 genotoxic effect on hFFs. **a**, SNP choice for scSNP-DNAseq analysis. Megabasic panel for detection of LOH after *ABRAXAS2* targeting in hFFs. Schema of 4 possible LOH profiles of the targeted chromosome 10 (no LOH = WT, tLOH I (LOH10qI, red allele), tLOH II (LOH10qII, blue allele) and 10qp). % of copy-neutral and copy-loss LOHs for experimental conditions with more than 5 cells by LOH profile. **b**, Cell cycle analysis in presence of AZD7648,

**Supplementary Fig 4:**
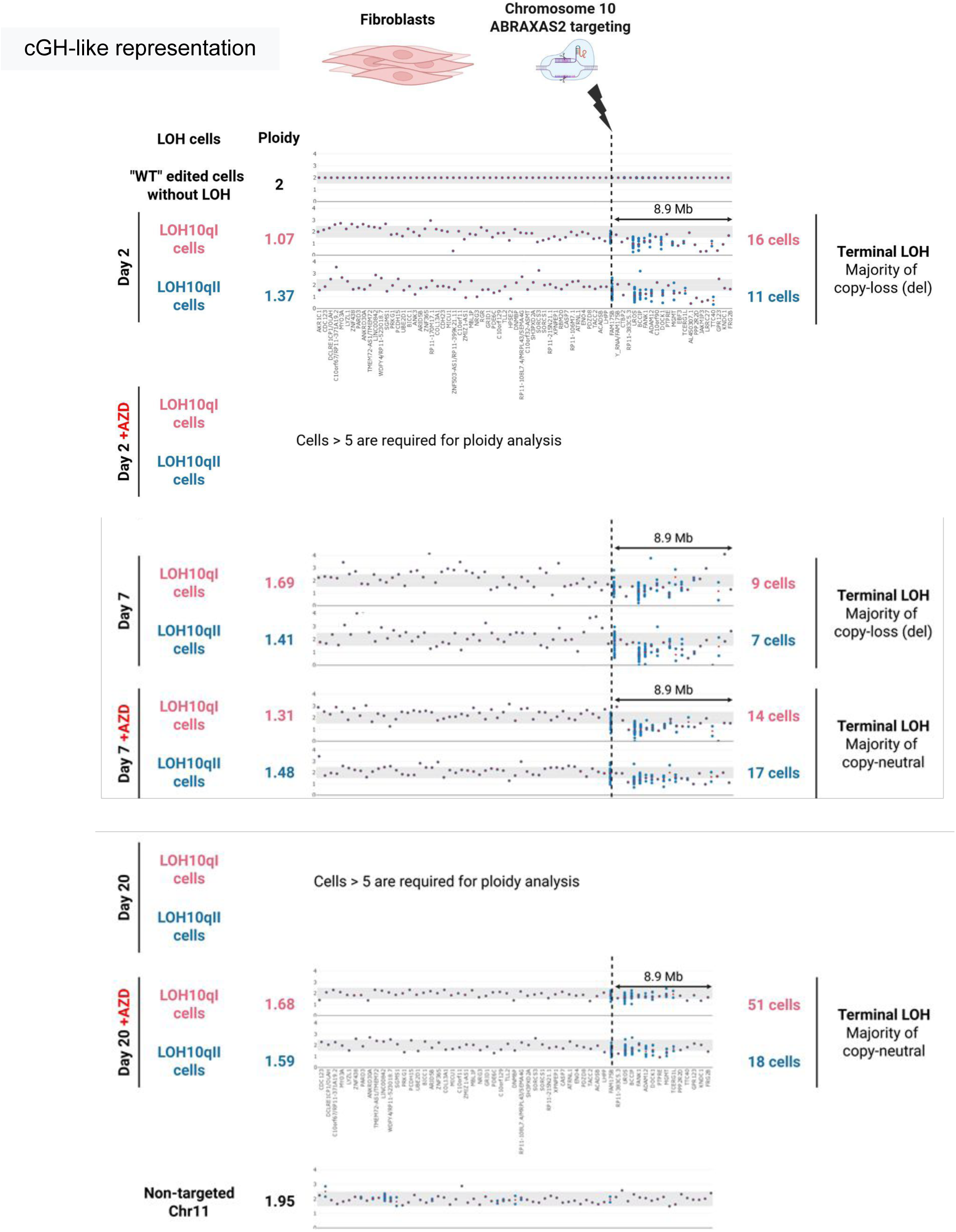
Ploidy analysis of edited cells with LOH. CNV and cGH-like representation of LOH^+^ hFF population after *ABRAXAS2* targeting in presence or absence of AZD7648 (upper panel, day 2; middle panel day 7; lower panel, day 20 and the non-targeted control chromosome 11).

**Supplementary Fig 5:**
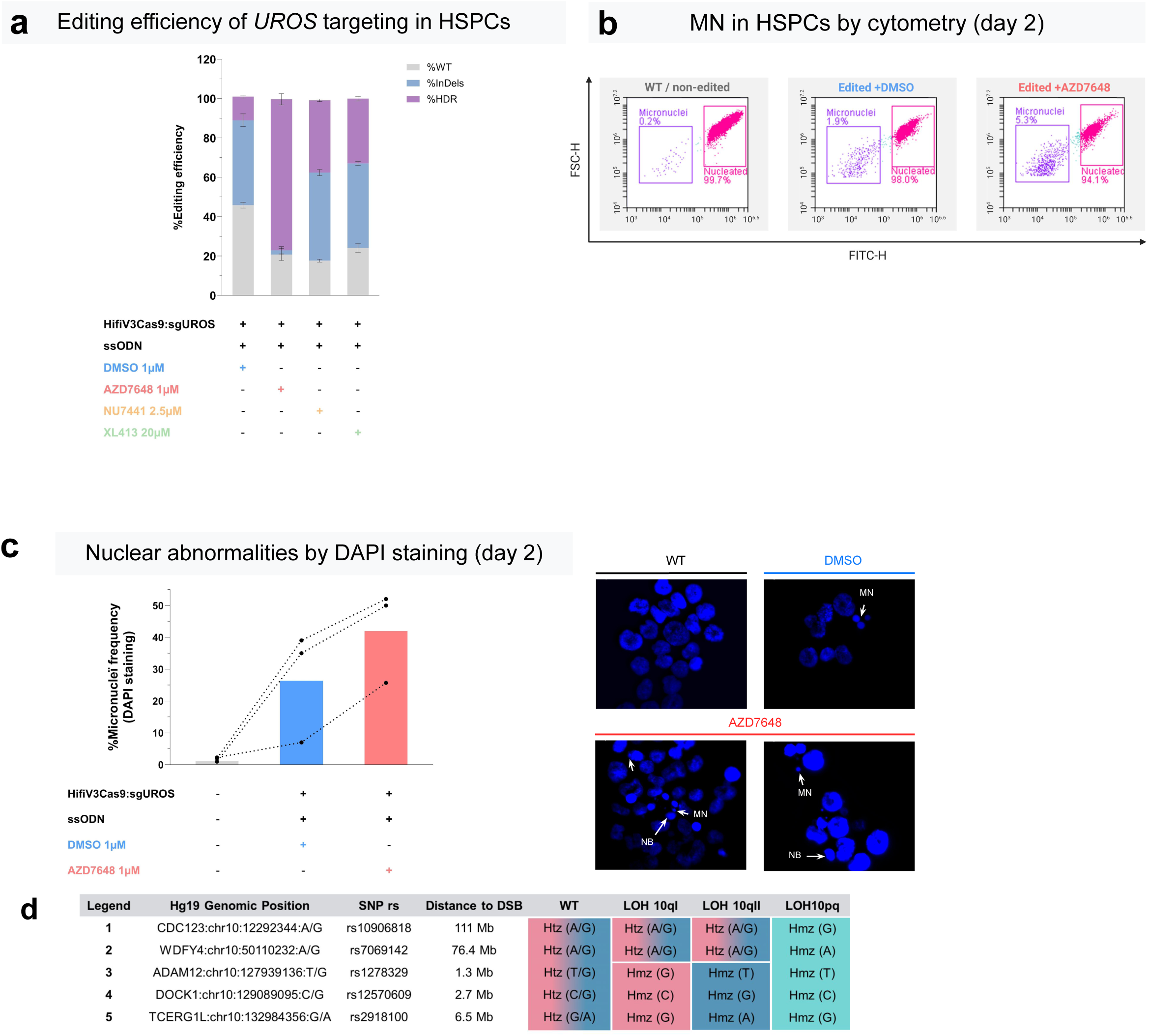
AZD7648 genotoxic effect in HSPCs. **a,** Editing efficacy of *UROS* editing, measured by PCR, Sanger sequencing and DECODR in presence of HDR booster, in HSPCs (Mean ± SD). **b**, illustrative cytometry dot plots of MN analysis in HSPCs (at least 10,000 cells per condition). **c**, **left**: fluorescence microscopy : % of nuclear abnormalities by DAPI staining at day 2 (n=3 independent experiments, n=500 cells). **right:** illustrative pictures of DAPI staining taken by Leica DM5500 B (Leica microsystem, Wetzlar, Germany) with the 63x objective lens (i.e. 630x magnification).**d,** SNP panel used for LOH detection in HSPCs (for scSNP-DNAseq). SNP profile of the different cell populations detected (WT, LOH 10qI, LOH 10qII, LOH 10qp). MN : micronuclei. NB: nuclear buds.

**Supplementary Fig 6:**
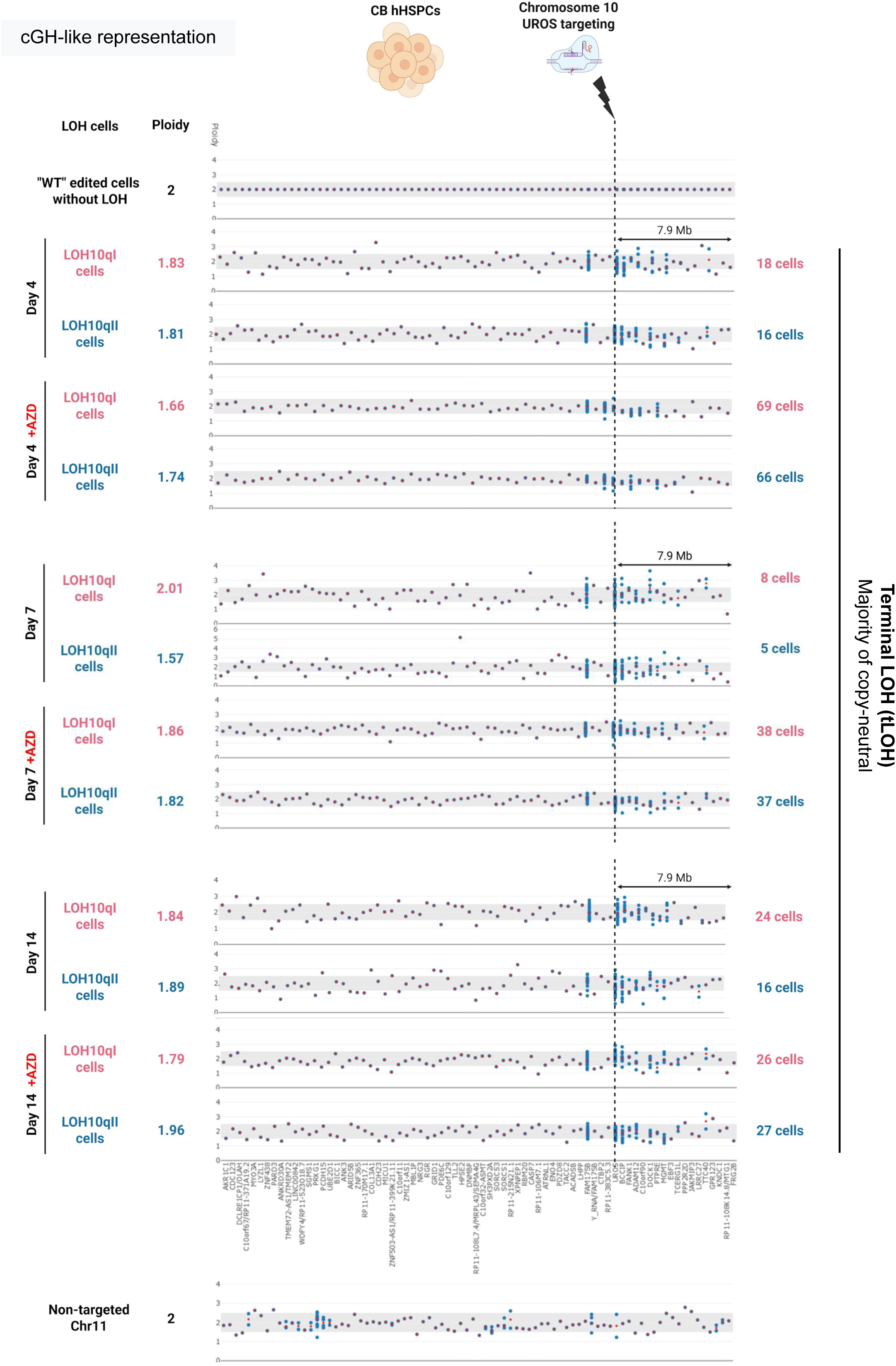
Ploidy analysis and cGH-like representation of LOH^+^HSPC population after *UROS* targeting in presence or absence of AZD7648 at day 4, day 7, day 14 of the targeted chromosome 10 and of the non-targeted control chromosome 11.

**Supplementary Fig 7:**
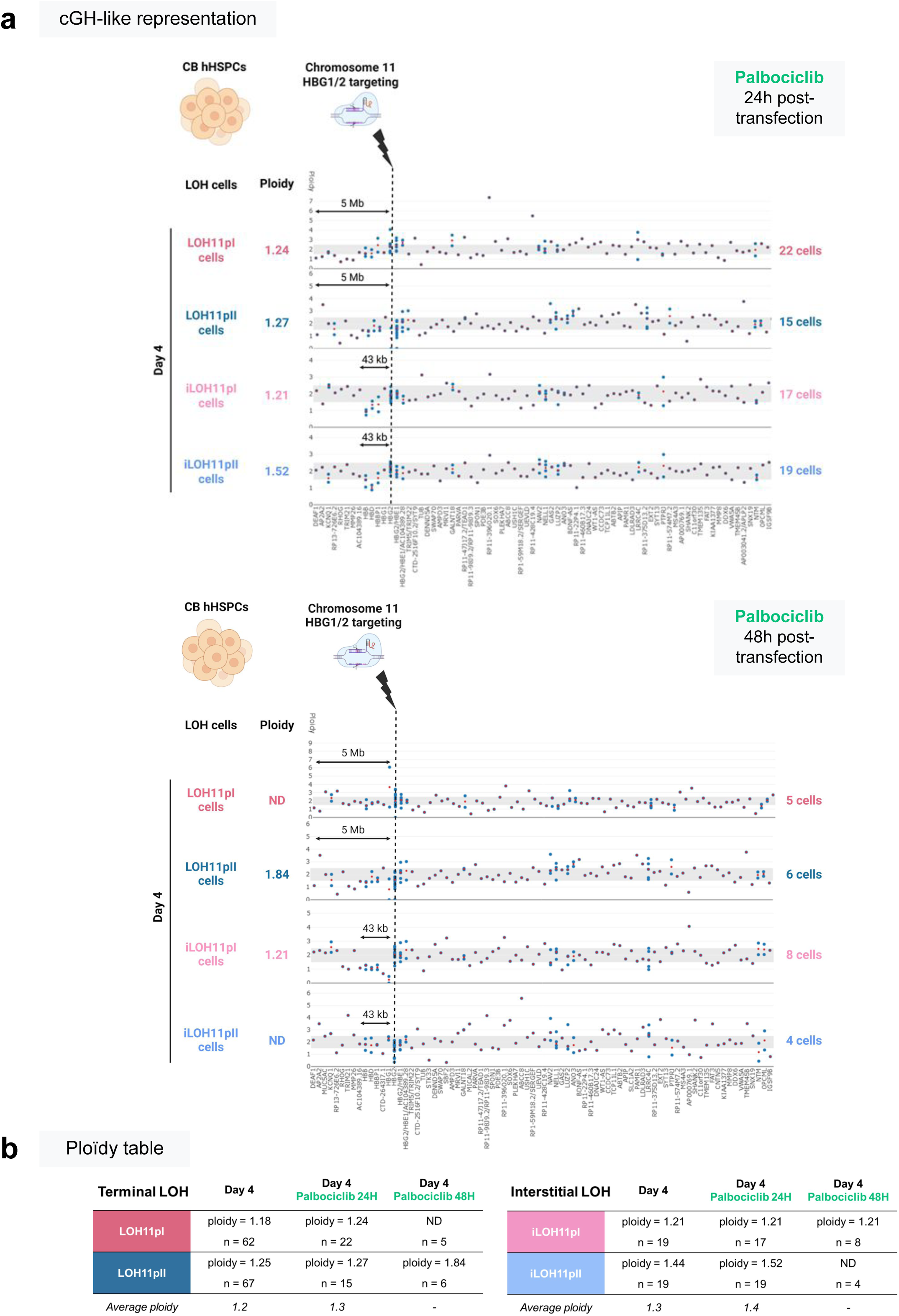
Ploidy analysis and cGH-like representation of LOH^+^ HSPC populations targeting *HBG1/2* promoters in presence of palbociclib (24h or 48 hours after transfection). **a,** Ploidy analysis and cGH-like representation of LOH^+^ HSPC populations after *HBG1/2p* targeting in presence of palbociclib (upper panel, palbociclib 24h; lower panel, palbociclib 48h). ND, non determined, ≤ 5 cells.**b,** Table of terminal and interstitial LOHs ploidies

**Supplementary Fig. 8:**
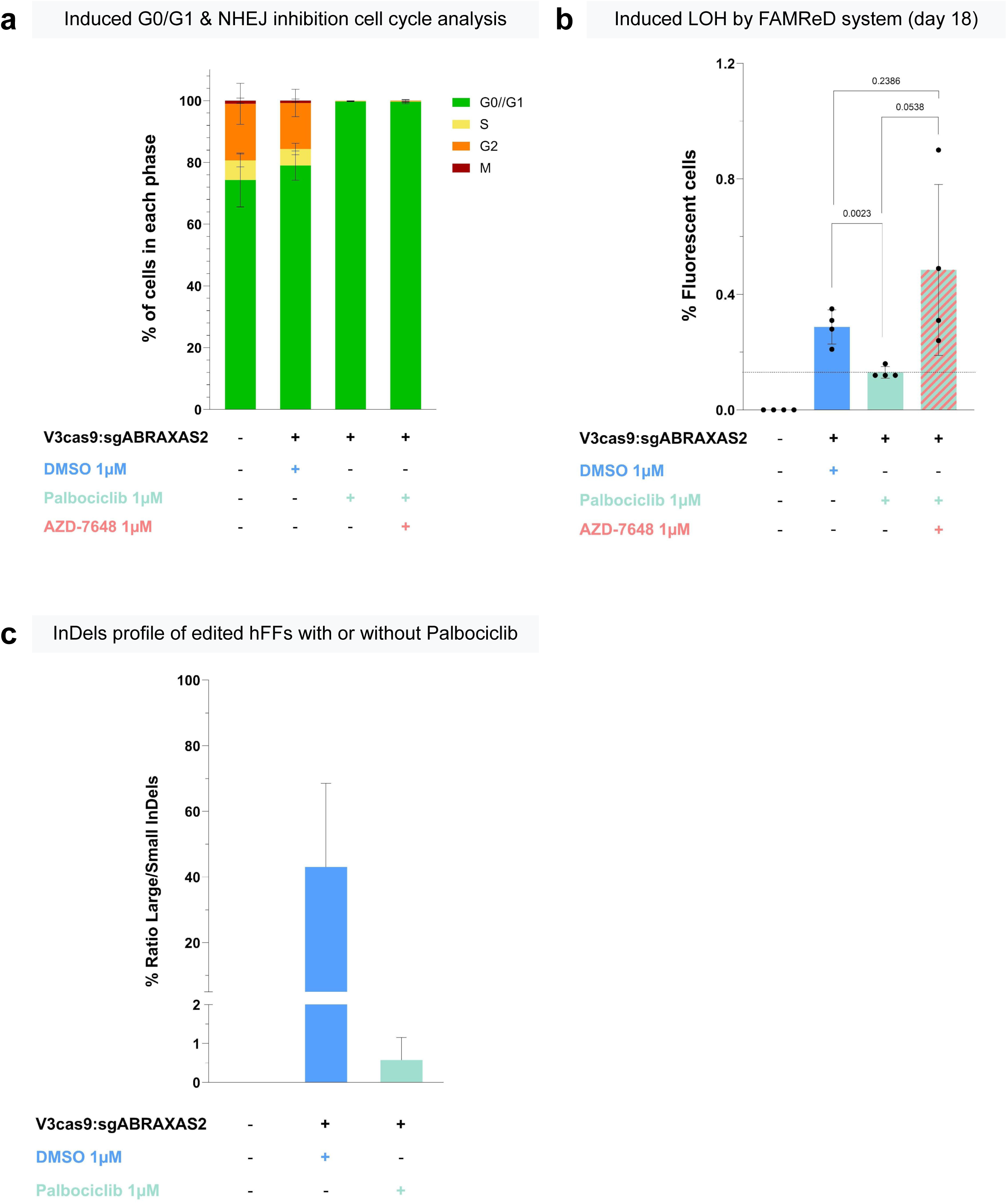
Mechanism of palbociclib protecting effect on genotoxicity by NHEJ. **a**, cell cycle analysis of edited hFFs in presence of palbociclib (1µM, 36h) with or without AZD7648 (1µM, 48h). **b**, % of cells with megabase scale LOH (fluorescent cells) after *ABRAXAS2* editing (by FAMReD system). Mean ± SD, n= 4 independent experiments, One-way ANOVA tests. **c**, Indels profiles of edited hFFs with or without palbociclib by sequencing and DECODR software. Mean ± SEM, n= 4 independent experiments. Small indels : <5bp from NHEJ. Large Indels: > ou = 5bp from MMEJ.

**Supplementary Fig. 9:**
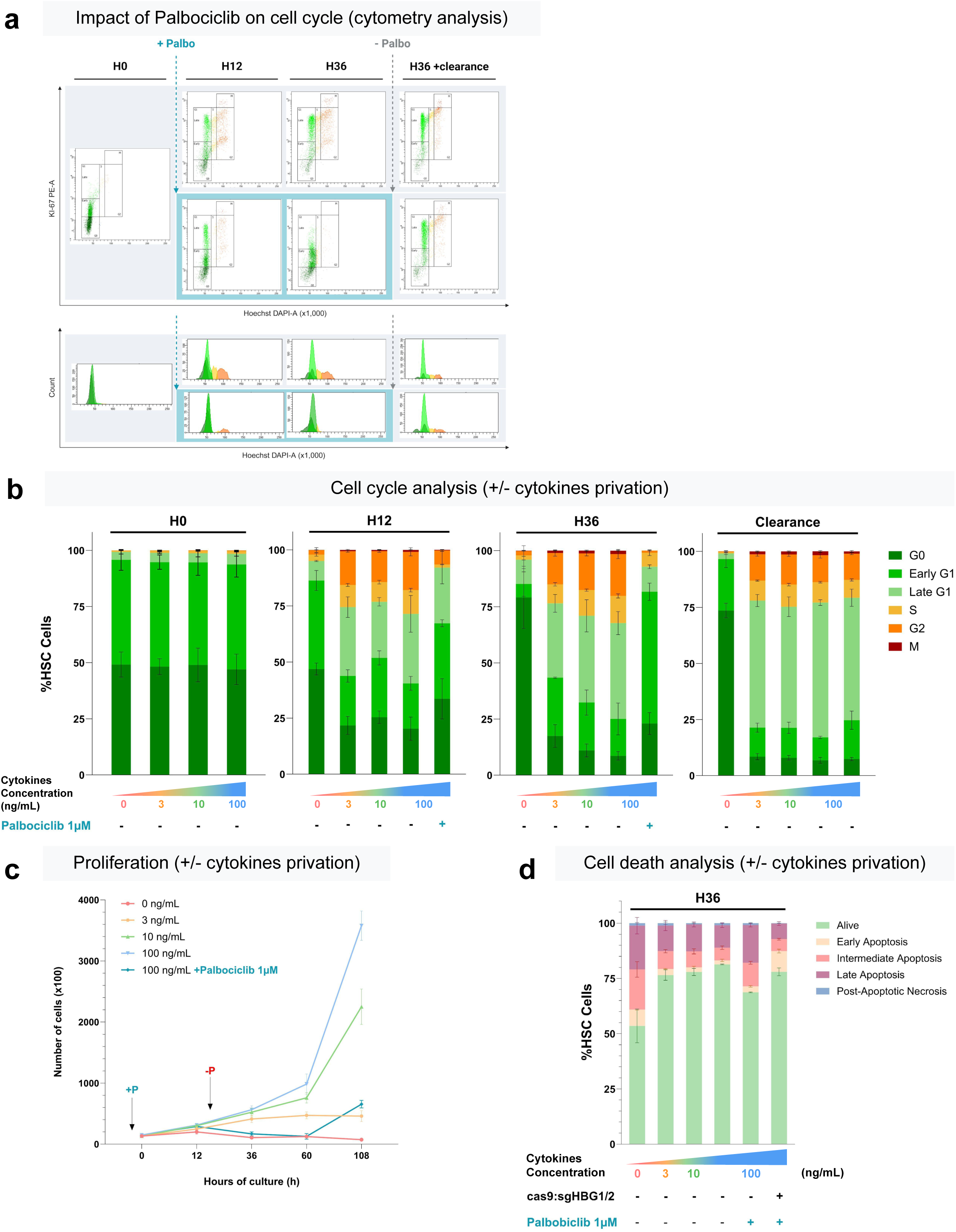
Palbociclib vs Cytokines privation. **a,** illustrative cytometry pictures of cell cycle analysis. HSPCs were analyzed at thawing (H0), 12h and 36h with palbociclib exposure and after a 48h-clearance. **b**, comparison of cell cycle phase repartition. (n= 3 independent experiments). **c**, proliferation assay during 108 hours by cell count. (n = 3 independent experiments). **d**, cell death analysis by annexin V and propidium iodure after 36h of HSPC culture with cytokine starvation (0-100ng/mL of SCF, TPO and FLT3L) or palbociclib exposure (n = 3 independent experiments). Palbociclib data from Fig 5b and 6b have been replotted to facilitate comparison to cytokine starvation.

**Supplementary Fig 10:**
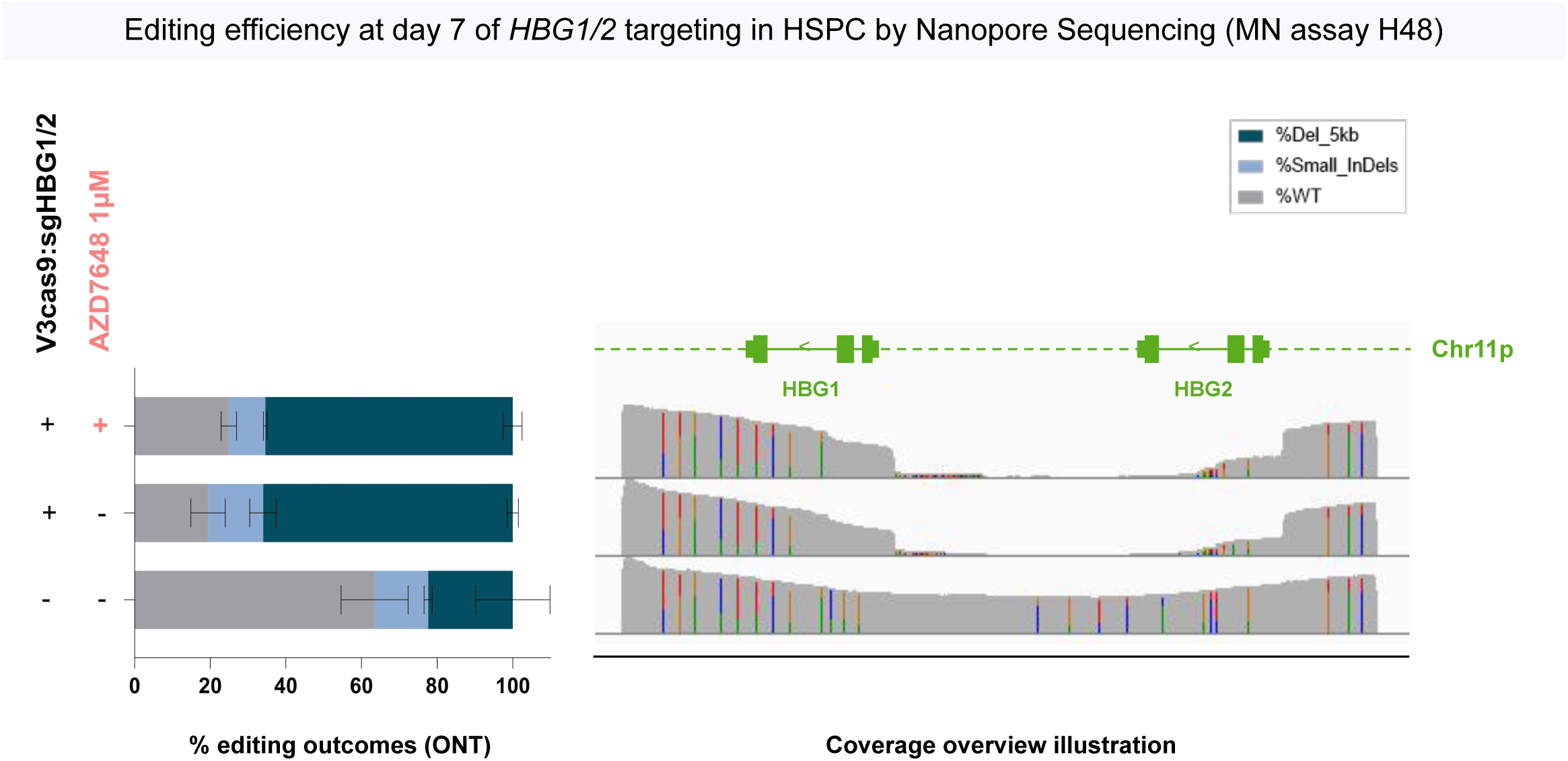
Editing efficiency at day 7 measured by long-range PCR (10 kb) and Nanopore sequencing for *HBG1/2* editing (MN assay H48). **Left panel**, histogram of genomic events after *HBG1/2* promoter targeting by CRISPR-Cas9 nuclease in HSPCs. Grey = WT sequence; light blue = indels, mutation or sequencing error; dark blue = 5Kb-deletion in HSPCs non edited, edited, and edited with palbociclib. **Right panel,** coverage overview illustration of NGS by Nanopore visualized with IGV.

**Supplementary Table 1:**
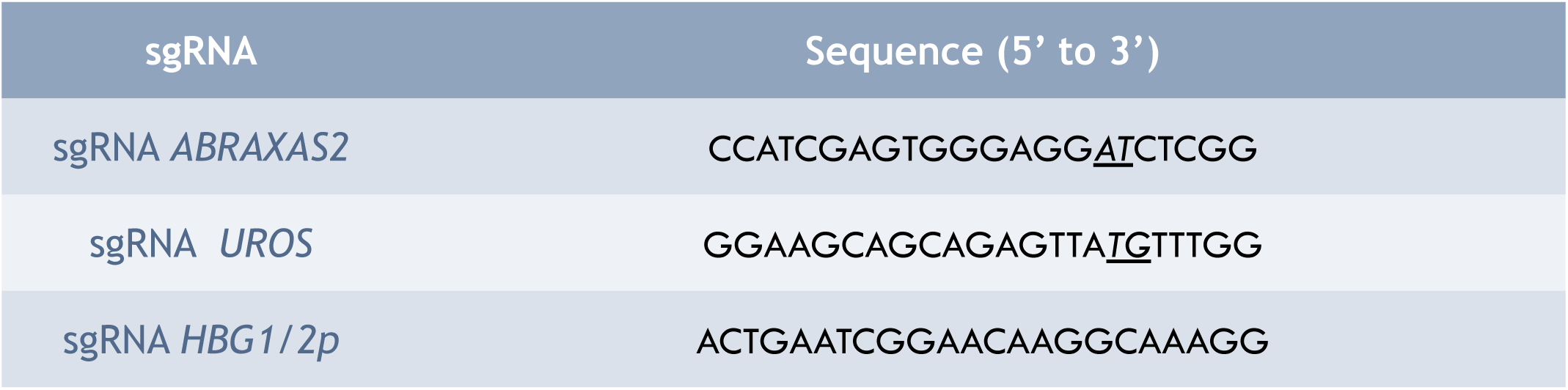
gRNA sequences for CRISPR-Cas9 editing.

**Supplementary Table 2:**
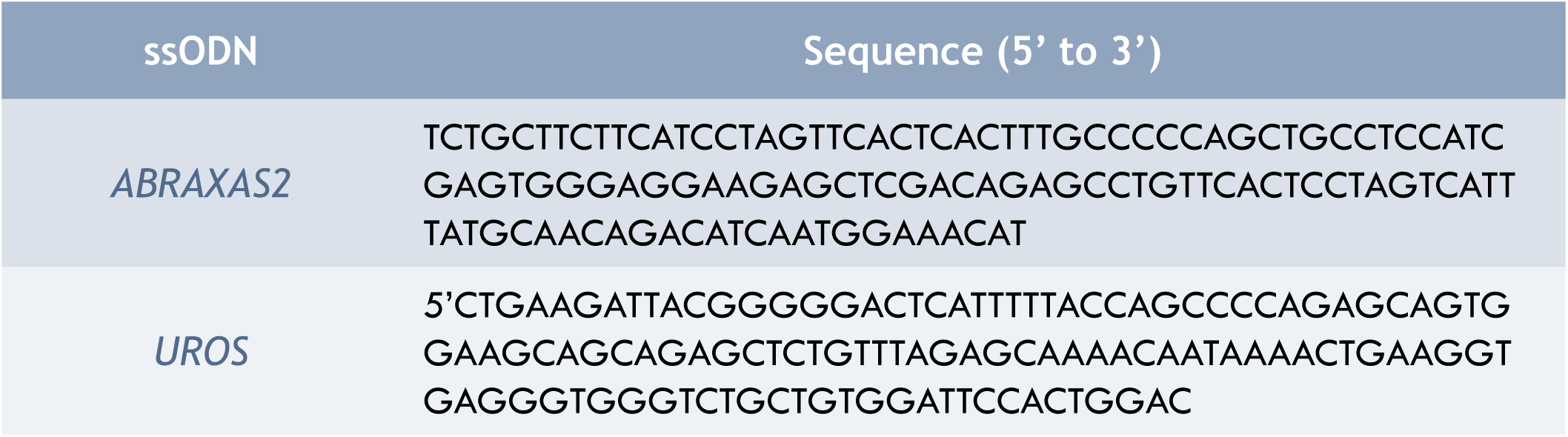
HDR ssODN template sequences for CRISPR-Cas9 HDR editing.

**Supplementary Table 3:**
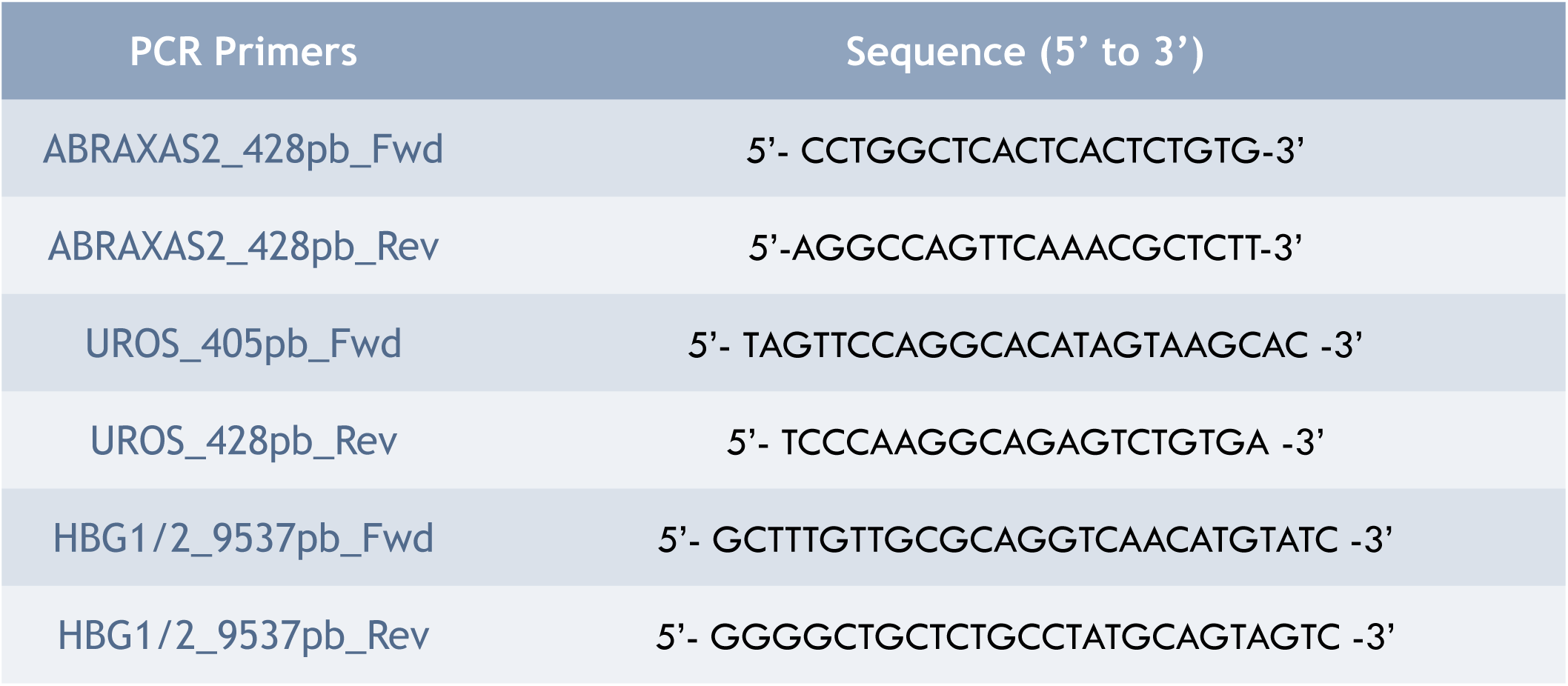
PCR primer sequences.

**Supplementary Table 4:** scDNAseq Global Metrics.

|  | Global |
| --- | --- |
| Amplicons (n) | 452 |
| Panel uniformity (average %) | 84,54 |
| Mean reads/cell/amplicon | 44,25 |
| Reads mapped to target (%) | 76,4 |
| Ado_rate (%) | 17,05 |
| Amplicon >10X (%) | 79,46 |

## Notes

### Summary of Updates

New co-first author (C Berges) New results : in vivo evidence that LOH events in HSPCs disappear after transplantation, mechanistic experiments demonstrating that protective efect of palbociclib is NHEJ-dependent, and functional studies showing improved engraftment of edited HSPCs

